# CRISPR screen identifies CNIH1 as a selective driver of GPCR export

**DOI:** 10.1101/2025.10.27.684930

**Authors:** Kevin Assoumou, Polina Drugachenok, Itziar Muneta Arrate, Xuefeng Zhang, Simon M. G. Braun, Miriam Stoeber

## Abstract

G protein-coupled receptors (GPCRs), the largest family of transmembrane proteins, transduce extracellular stimuli into intracellular signaling cascades to orchestrate human physiology. The transport of newly synthesized receptors from the endoplasmic reticulum (ER) to the plasma membrane (PM) determines cellular responsiveness to incoming ligands, yet the molecular machinery governing GPCR export remains incompletely defined. Here, we combine a synchronized cargo-release assay with a genome-wide CRISPR/Cas9 screen to systematically map regulators of GPCR ER-to-PM transport. Focusing on the δ-opioid receptor (DOR), a prototypical class A GPCR, we identify CNIH1 as a dedicated export factor. In the absence of CNIH1, DOR is retained intracellularly with immature glycosylation, and drives reduced PM signaling. CNIH1 localizes to both ER exit sites and the Golgi, promoting the anterograde transport of a subset of class A GPCRs. Opioid receptors directly interact with CNIH1 and require its putative COPII-binding site for export. Distinct from other human cornichon homologs, CNIH1 defines a selective GPCR-sorting receptor that couples GPCR biosynthesis to signaling competence.

## Introduction

The precise control of receptor abundance in the plasma membrane (PM) is essential for cells to accurately respond to extracellular signals. G protein-coupled receptors (GPCRs), the largest and most diverse family of membrane receptors, share a characteristic seven-transmembrane domain topology and mediate signaling by a wide array of endogenous and therapeutic ligands^1,2^. Cell surface GPCR levels reflect the combined influence of biosynthetic delivery, endocytosis, and recycling. Extensive work has defined the mechanisms that regulate ligand-dependent receptor internalization and endosomal sorting, often through live-cell imaging of labeled receptors and ligands^3,4^. By contrast, the molecular machinery that governs the anterograde transport of newly synthesized GPCRs through the secretory pathway remains poorly defined. This gap stems partly from the low abundance of nascent receptors at steady state and their inaccessibility to various labeling strategies. Real-time visualization and quantification of receptor passage through ER and Golgi are therefore crucial for defining the principles of GPCR surface delivery.

The anterograde transport of GPCRs employs a series of conserved synthesis, folding, and sorting steps shared with other membrane and secretory cargoes. These processes involve coordinated interactions with chaperones, vesicular coat proteins, and small GTPases, which are often determined by receptor-embedded motifs^5–8^. Beyond the canonical machinery, emerging studies have uncovered several factors that specifically tune the biosynthetic transport of individual GPCRs^9–12^. A systematic and unbiased identification of regulator proteins promises to uncover fundamental cellular principles that ensure the selectivity of GPCR transport and thereby signaling.

Opioid receptors, prototypical members of the large rhodopsin-like class A GPCR family, regulate pain perception and mood states, highlighting their physiological and therapeutic relevance^13,14^. The δ-opioid receptor (DOR), a biochemically well-characterized member of this family, undergoes folding and ER export as rate-limiting steps in its biosynthetic maturation^15,16^. At steady state, DOR localizes to both the PM and the Golgi apparatus, and activation of distinct subcellular receptor pools elicits divergent signaling outcomes^17–19^. Given the functional importance of DOR localization, we here focused on DOR as a model to dissect GPCR biosynthetic transport.

To follow nascent GPCRs with high temporal and spatial resolution, we employed the Retention Using Selective Hooks (RUSH) system^20^. RUSH enables the synchronized release of newly synthesized cargo from the ER, allowing quantitative measurement of trafficking kinetics under defined physiological and perturbative conditions. We combined the RUSH system for the first time with a genome-wide CRISPR-Cas9 knockout screen to systematically identify proteins that promote or restrict DOR cell surface transport. This integrated strategy enabled the unbiased discovery of regulators of GPCR export and revealed cornichon homolog 1 (CNIH1) as a key driver of anterograde transport. In mammals, the four CNIH proteins (CNIH1-4) have been implicated in the biosynthetic trafficking of specific clients, including AMPA-type ionotropic glutamate receptors^21,22^, but a role for CNIH1 in GPCR export has not been established. Here we found that loss of CNIH1 led to intracellular retention of biosynthetic DOR, which exhibited immature glycosylation and reduced PM signaling. CNIH1 localizes to both ER exit sites and the cis-medial Golgi, facilitating the anterograde transport of a subset of class A GPCRs, acting through a putative COPII-binding motif and direct receptor interaction. Together, these findings establish CNIH1 as a dedicated GPCR-sorting receptor, functionally distinct from other human CNIH proteins, and reveal a new mechanism of cargo selectivity within the early secretory pathway that affects GPCR signaling fidelity.

## Results

### GPCR anterograde trafficking is regulated

To investigate GPCR ER-to-PM transport, we adapted the synchronized RUSH export assay for diverse receptors of interest. In our RUSH setup, the GPCR of interest was fused at the N-terminus with the streptavidin-binding peptide (SBP) and GFP, and co-expressed with a streptavidin-KDEL hook protein consisting of streptavidin fused to the ER retention signal KDEL (**Fig. 1a**)^20^. In the absence of biotin, the GPCR is retained in the ER due to interaction of SBP with streptavidin-KDEL in the organelle lumen. Upon biotin addition, SBP is displaced from streptavidin, releasing the receptor for synchronized trafficking that can be visualized via GFP (**Fig. 1a**). Using the PiggyBac transposon approach^23,24^, we generated polyclonal HeLa cells stably expressing both the GPCR and the ER hook from a bicistronic construct **(Extended Data Fig. 1a)**.

**Figure 1.**
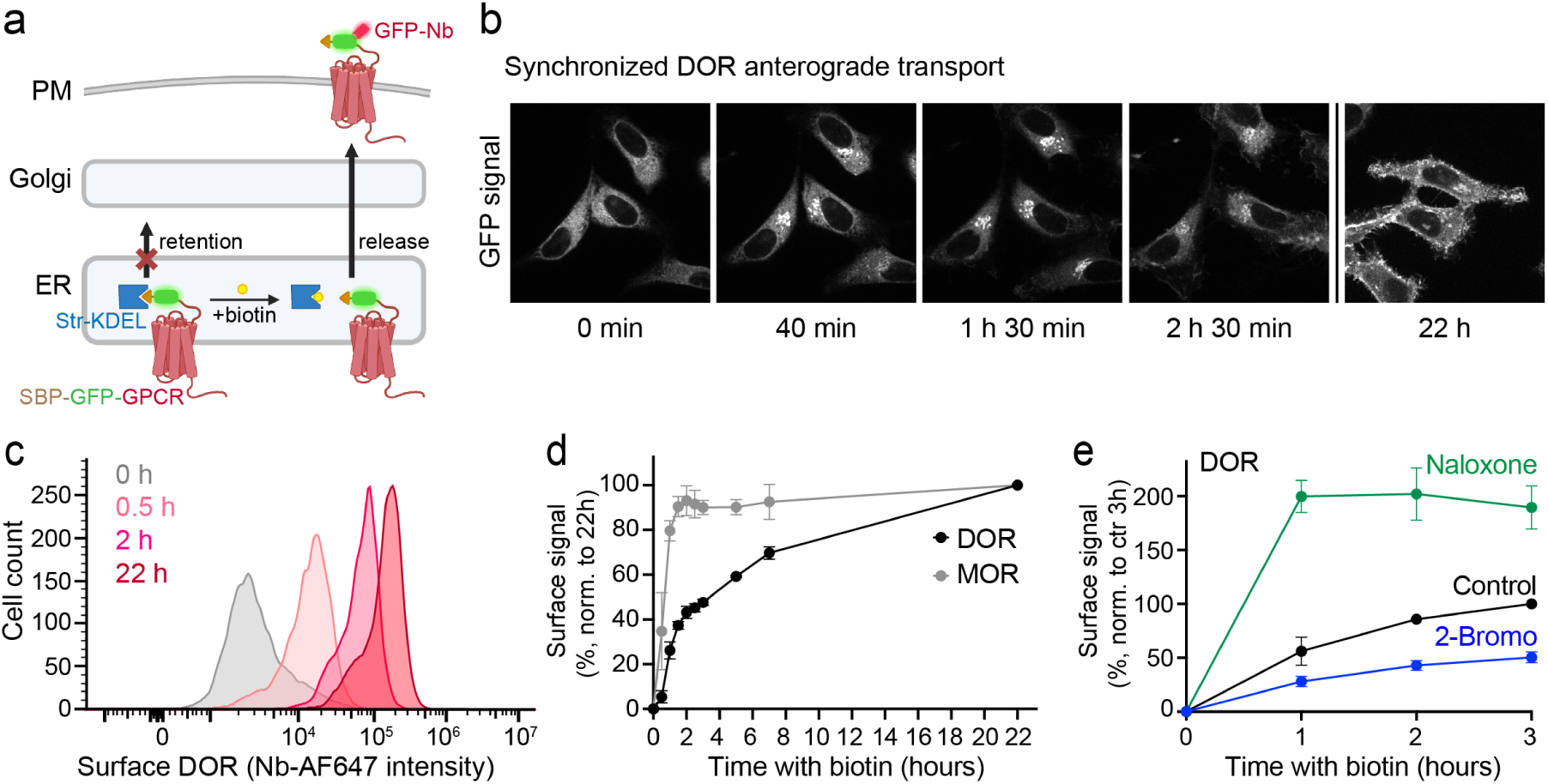
Regulated GPCR anterograde transport detected with the RUSH assay. **(a)** Schematic representation of the RUSH assay. Left: retention in the absence of biotin. Right: release of GPCRs from the ER. Str-KDEL: streptavidin-fused ER hook protein; SBP: streptavidin-binding peptide. **(b)** Confocal time series of DOR-RUSH HeLa cells imaged at indicated time points after biotin addition (80 μM). At 22h, different cells are shown. Images are single confocal slices. Scale bar, 10 μm. **(c)** Flow cytometry histograms of DOR-RUSH HeLa cells stained with AF647-labeled GFP-Nb at indicated times after biotin addition. **(d)** Kinetics of PM arrival of DOR and MOR based on flow cytometry analyses. The AF647 signal at each time point was normalized to the 22 h signal (set to 100%) for each receptor. Data are mean ± SD from 3 independent experiments. **(e)** Kinetics of DOR PM targeting in control cells and in cells pre-treated with 10 μM naloxone (10 min pre-treatment) or 100 μM 2-bromopalmitate (2-Br, 16 h pre-treatment) prior to biotin addition. Signal in control cells at 3 h was set to 100%. Data are mean ± SD from 3 independent experiments.

We focused our initial analyses on the DOR that was homogeneously expressed and showed low receptor surface levels in the absence of biotin **(Extended Data Fig. 1b,c**). Live-cell imaging by confocal fluorescence microscopy revealed efficient ER retention in the absence of biotin (**Fig. 1b**). Following biotin addition, DOR relocalized and exhibited a prominent pool in the Golgi between 40 and 90 min, which was followed by onward transport to the PM (**Fig. 1b**). To quantify DOR cell surface arrival over time, we incubated cells with biotin for different durations, stained cells with a fluorescently conjugated anti-GFP nanobody (GFP-Nb)^25^, and measured the signal by flow cytometry. DOR levels in the PM gradually rose up to a 25-fold increase at 22 hours, while total receptor levels monitored via GFP remained constant during the time course (**Fig. 1c,d, Extended Data Fig. 1b-d**).

We next quantified the anterograde transport of the highly homologous mu-opioid receptor (MOR), which was expressed at similar levels to DOR in the stable RUSH HeLa cells **(Extended Data Fig. 1e)**. Confocal microscopy and flow cytometry analyses confirmed efficient ER retention of MOR and the biotin-induced synchronized transport to the PM **(Extended Data Fig. 1d,f,g)**. Strikingly, the export kinetics of MOR and DOR strongly differed. Within 2 hours, MOR reached maximal PM levels that remained constant thereafter (export half time t1/2 = 45 min), while DOR ER-to-PM trafficking was markedly slower (t1/2 = 3.5 h) and showed maximum surface levels at 22 h (**Fig. 1d**). The kinetics differences were independent of the receptor expression level, as cells gated for low or high MOR or DOR levels showed a similar export behavior **(Extended Data Fig. 1h,i**).

Because a C-terminal motif in DOR has been implicated in regulating anterograde trafficking in neurons^26,27^, we tested whether this region contributes to the distinct kinetics of DOR and MOR surface arrival. We generated a chimeric receptor in which the MOR C-tail was replaced by that of DOR (MOR-DOR). In stable RUSH HeLa cells, MOR-DOR exhibited significantly slower biotin-induced surface targeting compared with MOR **(Extended Data Fig. 1j,k**). The findings support a role of the DOR C-terminus in defining receptor-specific anterograde transit and demonstrates that the RUSH assay identifies differences in the transport of individual GPCR cargos.

We next asked whether the assay could detect altered trafficking under pharmacological or biochemical perturbations. For this, we preincubated DOR-RUSH cells with naloxone, a permeant antagonist of DOR that acts as a pharmacological chaperone for receptors in the ER, and has been shown to enhance surface targeting of biosynthetic DOR^16,28^. Indeed, naloxone incubation prior to biotin addition produced a 2-fold increase in DOR surface levels at 3 hours after release (**Fig. 1e**). The enhanced PM targeting was dependent on the naloxone concentration (EC50 = 100 nM) **(Extended Data Fig. 2a)**. MOR, which also binds naloxone with high potency, showed a similar increase in receptor surface arrival in the presence of naloxone **(Extended Data Fig. 2b)**. We also detected up to 1.7-fold higher PM levels of the serotonin 5-HT2A receptor (5-HT2AR) when 5-HT2AR-RUSH cells were pre-incubated with the permeant antagonist ketanserin before adding biotin **(Extended Data Fig. 2c)**, showing that pharmacological chaperones can promote the export of diverse biosynthetic GPCRs. Importantly, the results demonstrated that the synchronized GPCR anterograde transport assay was not operating at maximum export capacity in unperturbed cells.

Finally, we identified conditions that impaired passage of nascent GPCRs through the secretory pathway. Preincubation of DOR-or MOR-RUSH cells with 2-bromopalmitate, a palmitoylation inhibitor, decreased the surface targeting of both receptors by ∼50% (**Fig. 1e, Extended Data Fig. 2b**), consistent with previous studies linking opioid receptor palmitoylation to surface abundance^29^.

Together, the results establish the synchronized RUSH release assay as a quantitative and sensitive approach to monitor GPCR-specific ER-to-PM trafficking in intact cells, and robustly detects conditions with altered anterograde transport behavior. Importantly, the assay was amenable to high throughput screening techniques such as FACS, enabling the analyses of GPCR transport in individual cells, which we took advantage of in our subsequent screen to identify proteins controlling DOR export.

### Genome-wide CRISPR screen identifies regulators of DOR anterograde transport

To identify cellular factors that regulate GPCR ER-to-PM transport, we combined the quantitative RUSH assay with a genome-wide CRISPR/Cas9 knockout (KO) screen (**Fig. 2a**). We focused on DOR because its comparatively slow and inefficient PM targeting (**Fig. 1d**) provided a sensitive dynamic range for detecting both enhancers and inhibitors of GPCR anterograde transport in the same experimental workflow.

**Figure 2.**
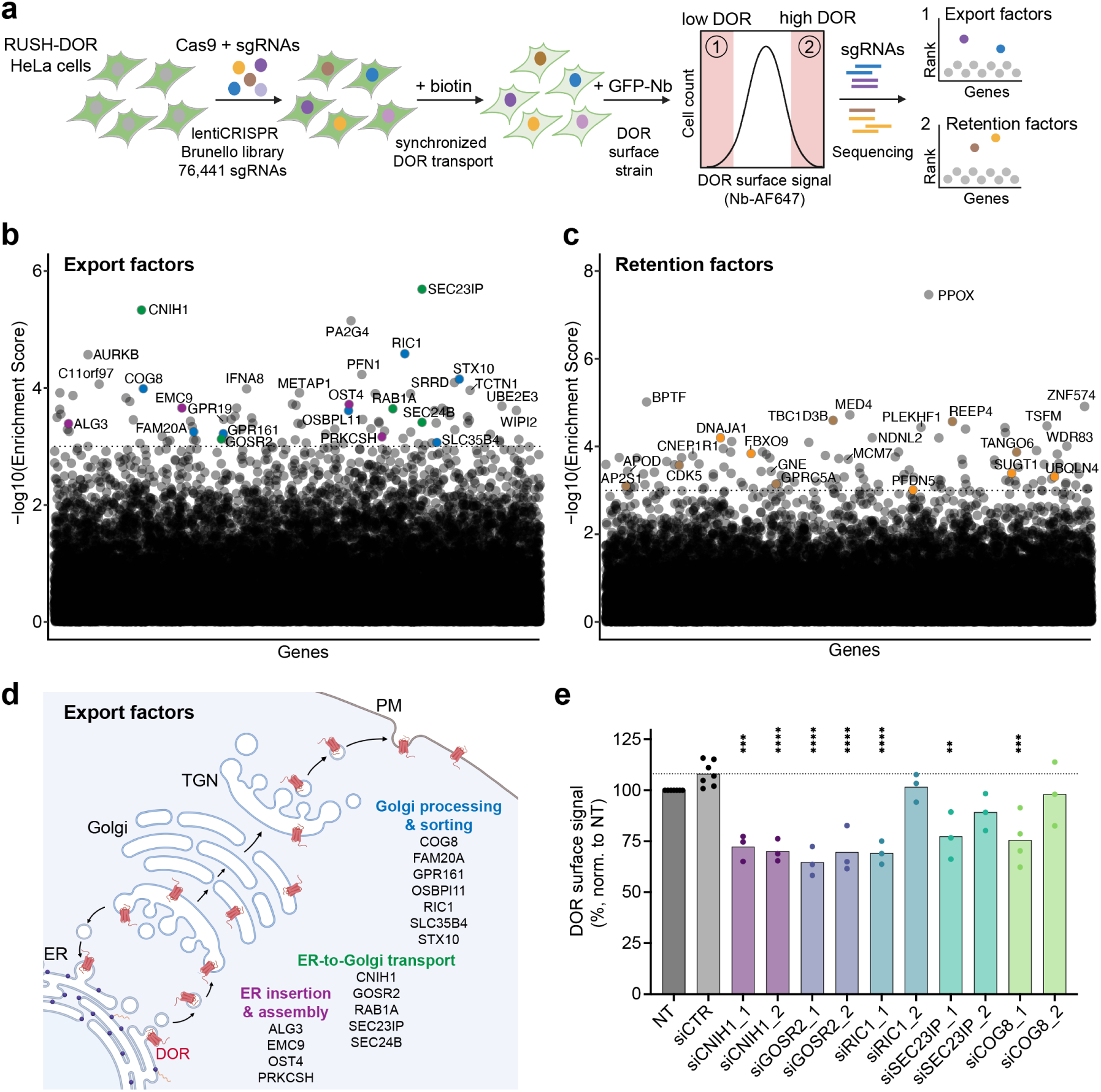
Genome-wide CRISPR/Cas9 screen identifies regulators of DOR anterograde transport. **(a)** Workflow of the genome-wide CRISPR/Cas9 KO screen. DOR-RUSH HeLa cells were transduced with the lentiviral Brunello sgRNA library and selected with puromycin. On the assay day, biotin was added for 1.5 h, and cells with the 10% lowest and highest DOR surface levels, as well as the unsorted population, collected for genomic DNA extraction and sequencing to determine enriched sgRNAs. **(b, c)** Plot of gene enrichment scores from MAGeCK analysis of sgRNA counts with a cut-off of 0.001 (dotted line) of export factors (b) or retention factors (c). Each gene targeted by the sgRNA library is indicated as a black dot. Factors belonging to functional categories of interest are color-coded. Export factors: purple = ER insertion and assembly, green = ER-to-Golgi transport, blue = Golgi processing and sorting. Retention factors: brown = membrane trafficking, orange = protein folding and degradation. All gene enrichment scores are provided in Extended Data Table 1. **(d)** Schematic of the GPCR anterograde trafficking pathway with identified export regulators that likely act at distinct steps. **(e)** Validation of screen hits by siRNA-mediated knockdown in DOR-RUSH HeLa cells, followed by quantification of DOR export using flow cytometry. DOR export at 1.5 h was normalized to non-treated (NT) control cells (set to 100%). n = 3 independent experiments. Significance between non-targeting control siRNA (siCTR) and targeted siRNAs determined by ordinary one-way ANOVA with Tukey’s multiple comparisons test; *, P < 0.05; **, P < 0.01; ***P < 0.001.

We generated KO cells of single genes in the DOR-RUSH cell line by transducing the cells at low multiplicity of infection (MOI of 0.3) with the human CRISPR KO pooled lentivirus Brunello library^30,31^. The library consists of 3-4 single guide RNAs (sgRNAs) for each of the 19,114 targeted human genes as well as control sgRNAs. To maintain library diversity, we used a coverage of 500x per sgRNA. Cells with stably integrated spCas9 and respective sgRNA were selected by puromycin (**Fig. 2a**).

To screen for DOR anterograde transport regulators, the pooled KO cells were incubated with biotin for 1.5 h, a time point at which DOR export was particularly sensitive to modulation (**Fig. 1e**) and the possible influence of constitutive endocytosis was likely minimal. Surface-localized DOR was labeled with an Alexa Fluor 647-conjugated GFP-Nb, and cells were sorted by flow cytometry into low and high surface DOR populations corresponding to the lowest and highest 10% of Nb-AF647 signals (**Fig. 2a**). The unsorted population served as the control. Each cell population was processed for next generation sequencing to identify the enriched sgRNAs (**Fig. 2a**). We detected >97% of all individual sgRNAs in the control population and the vast majority of sgRNAs had read counts between 100 to 1000 **(Extended Data Fig. 3a)**, confirming the diverse representation and anticipated sgRNA coverage in the screen.

MAGeCK analyses^32^ identified enriched sgRNAs by comparing the cell populations with low and high DOR surface levels. Applying a stringent enrichment score cutoff of 0.001, we identified 89 export factors (KO reduces DOR surface levels) (**Fig. 2b, Extended DataTable 1**) and 115 retention factors (KO increases DOR surface levels) (**Fig. 2c, Extended Data Table 1**). Gene ontology (GO) term enrichment analyses indicated that main functional categories associated with the export factors were intracellular protein transport, ER to Golgi vesicle-mediated transport, and protein N-linked glycosylation **(Extended Data Fig. 3d)**.

We further examined the hits based on their subcellular location of function (**Fig. 2d**). We identified ER folding and assembly factors such as the ER membrane insertase component EMC9, as well as enzymes involved in N-linked glycosylation (ALG3, OST4, and PRKCSH). ER-to-Golgi transport proteins included COPII complex proteins SEC24B and SEC23IP, the cornichon family protein CNIH1, the Rab GTPase RAB1A, and the Golgi SNARE GOSR2. We also identified Golgi processing and sorting factors, including a subunit of the conserved oligomeric Golgi complex COG8, the guanine nucleotide exchange factor RIC1, and the SNARE STX10, amongst others (**Fig. 2d**). These genes exhibited the highest enrichment scores (**Fig. 2b**) and top ranked p-values **(Extended Data Fig. 3b)**, and in most cases, all 4 individual sgRNAs targeting the respective genes were enriched **(Extended Data Fig. 3e)**.

Among the retention factors, we identified hit genes related to membrane trafficking, including for example the ER protein REEP4, the putative Rab GTPase-activating protein TBC1D3, non-cell cycle regulating cyclin-dependent kinase CDK5, the orphan GPCR GPRC5A, and the transport and Golgi organization protein TANGO6 (**Fig. 2c**). Additionally, we identified hit genes involved in protein folding and degradation as retention factors, such as (co-)chaperones DNJA1 and PFDN5, the E3 ubiquitin ligase component FBXO9, the SCF ubiquitin ligase component SUGT1, and the ubiquilin UBQLN4 (**Fig. 2c**). These genes also ranked among the top retention factors based on p-value **(Extended Data Fig. 3c)**.

In our follow up studies, we focused on the identified export factors and aimed to validate several top hits not previously known to promote DOR or GPCR anterograde transport. We downregulated the expression of five hit proteins using two independent siRNAs in HeLa DOR-RUSH cells and quantified DOR PM delivery by flow cytometry. Consistent with the CRISPR screen results, both siRNAs targeting CNIH1 or GOSR2 significantly reduced DOR ER-to-PM transport (**Fig. 2e**). DOR surface levels were also significantly reduced by one of the two siRNAs targeting RIC1, SEC23IP, or COG8. RT-qPCR confirmed efficient target mRNA knockdown **(Extended Data Fig. 4a)**, and total DOR levels, assessed via GFP fluorescence, remained unchanged except for a mild reduction upon COG8 depletion **(Extended Data Fig. 4b)**. This confirmed a role of the identified proteins in driving PM targeting of nascent DOR rather than total DOR abundance. Together, the flow cytometry-based RUSH CRISPR screen robustly identified previously unknown regulators of DOR anterograde transport for further characterization.

### CNIH1 promotes ER-to-Golgi transport, maturation, and PM signaling of DOR

One of the strongest hits from our screen and validation experiments was CNIH1, a member of the cornichon homolog (CNIH) protein family with four members in humans (CNIH1-4). The CNIH family is conserved in eukaryotes with the first described member, *Drosophila* cornichon Cni, required for delivery of TGFα from the ER to the oocyte surface during oogenesis^33,34^. The best characterized CNIH homolog is the *S. cerevisiae* protein Erv14p, which localizes to the ER and Golgi apparatus and promotes cargo incorporation into COPII vesicles by interacting with both cargo and COPII coat subunits^35–37^. In mammals, CNIH2 and CNIH3 regulate AMPA receptor trafficking^38,39^, and CNIH4 affects the export of the chemokine receptor CCR5, β2-adrenergic receptor, and the viral surface protein VSVG^11,40^. However, the function of CNIH1 has remained largely unknown. We therefore set out to define the role of CNIH1 in regulating DOR trafficking. We first examined the effect of CNIH1 loss on DOR transit through the secretory pathway. To deplete CNIH1, we used siRNA-induced knockdown or CRISPR/Cas9-mediated KO in DOR-RUSH HeLa cells. Both approaches strongly reduced CNIH1 expression, with undetectable protein levels in CNIH1 KO cells and a ∼70% decrease upon CNIH1 siRNA treatment, as detected by western blot (**Fig. 3a**). Total DOR expression was unchanged **(Extended Data Fig. 4c,d**). Cells depleted of CNIH1 showed reduced DOR surface targeting, which remained significantly lower at later time points after biotin addition when receptor surface levels plateaued (21h and 24h) (**Fig. 3b&c)**. Thus, loss of CNIH1 causes not only a kinetic delay in DOR PM delivery but also a sustained reduction in steady-state DOR surface levels.

**Figure 3.**
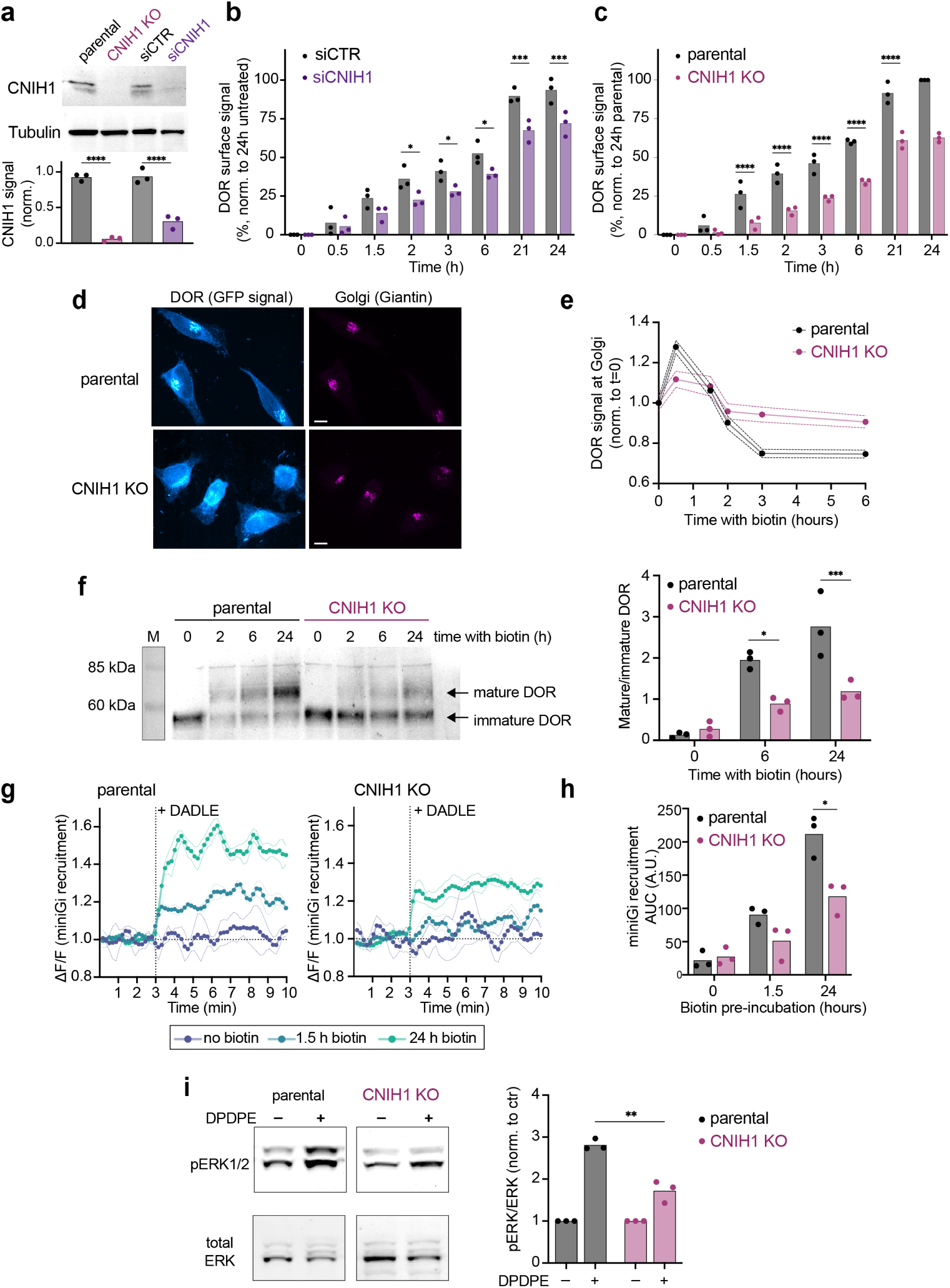
Loss of CNIH1 leads to intracellular DOR retention, immature glycosylation, and reduced signaling. **(a)** Western blot analysis of CNIH1 protein levels in parental and CNIH1 KO DOR-RUSH cells, and in cells treated with control (siCTR) or CNIH1-targeting siRNA (siCNIH1). CNIH1 signals were normalized to tubulin. n = 3 independent experiments. Significance was determined by one-way ANOVA and Tukey’s multiple comparisons tests; ****, P < 0.0001. **(b, c)** DOR PM levels at various time points after biotin addition in DOR-RUSH HeLa cells depleted of CNIH1 by siRNA (b) or CRISPR/Cas KO (c) shown relative to their respective controls. Surface DOR was quantified by flow cytometry normalized to the signal of parental DOR-RUSH cells at 24 h. n = 3 independent experiments. Significance was determined by two-way ANOVA with Šídák’s multiple comparisons tests; *, P < 0.05; ***, P < 0.001; ****, P < 0.0001. (**d**) Representative confocal microscopy images of parental and CNIH1 KO DOR-RUSH HeLa cells fixed 0.5 h after biotin addition. DOR (GFP signal) and the Golgi compartment (anti-Giantin immunolabeling) are shown. Images are maximum intensity Z-projections of confocal stacks. Scale bar, 10 μm. (**e**) Quantification of DOR fluorescence in the Golgi region in parental and CNIH1 KO DOR-RUSH cells after biotin addition. Cells were fixed and immunolabeled with anti-Giantin as in (d). Data are mean ± SEM error bands from 3 independent experiments. (**f**) Western blot analysis of DOR glycosylation 0, 2, 6 and 24 hours after biotin addition in parental and CNIH1 KO DOR-RUSH HeLa cells. DOR was isolated from cell lysates via GFP-Trap beads and treated with EndoH. Left panel: representative blot (anti-GFP). Right panel: Quantification of mature vs. immature receptor signals. n = 3 independent experiments. Significance was determined by two-way ANOVA with Šídák’s multiple comparisons tests; *, P < 0.05; **, P < 0.01; ***, P < 0.001. (**g**) Split-NanoLuc complementation assay measuring mGi1 recruitment to DOR in parental and CNIH1 KO DOR-RUSH cells expressing mGi1-LgBit and lyn11-SmBit. Cells were pre-incubated with biotin for 1.5 h or 24 h, or left untreated, and then stimulated with peptide agonist DADLE (100 nM, added at 3 min during acquisition). Graphs show mGi1 recruitment signal over time. Data are mean ± SEM error bands from 3 independent experiments. (**h**) Quantification of DADLE-induced mGi recruitment to DOR based on area under the curve (AUC) analysis in parental and CNIH1 KO cells as in (g). n = 3 independent experiments. Significance was determined by two-way ANOVA with Šídák’s multiple comparisons; *, P < 0.05. (**i**) Left: Western blot showing total ERK and phosphorylated ERK (pERK) levels in parental and CNIH1 KO DOR-RUSH cells after 5 min stimulation with 10 μM DPDPE (+) or vehicle (-). Right: Quantification of pERK/ERK ratios normalized to unstimulated control cells. n = 3 independent experiments. Significance was determined by two-tailed paired t-test; **, P < 0.01.

To more precisely characterize what step of DOR anterograde transport was affected by CNIH1 depletion, we performed immunofluorescence and monitored DOR passage from the ER to the Golgi apparatus, staining the Golgi compartment with a Giantin antibody (**Fig. 3d**). Upon biotin addition, we quantified the DOR signal in the Golgi area of control and CNIH1 KO cells. In cells devoid of CNIH1, Golgi-localized DOR was significantly reduced at early time points (0.5 h), while levels remained elevated at later time points (**Fig. 3d,e**). This was consistent with a delayed and less efficient ER-to-Golgi transport of biosynthetic DOR in the absence of CNIH1.

Previous work has shown that the transit of biosynthetic DOR through the Golgi apparatus is associated with the conversion of a precursor DOR form to a fully mature DOR glycoprotein containing N- and O-linked glycans^15^. The mature, glycosylated DOR migrates with a ∼10 kDa higher MW and is resistant to treatment with endoglycosidase H (Endo H), which selectively removes unprocessed high-mannose oligosaccharides but not fully processed glycans. Since we detected inefficient Golgi targeting in the absence of CNIH1, we next tested whether DOR conversion into the mature, Endo H-resistant glycoprotein was impaired. In DOR-RUSH control cells, we observed efficient, time-dependent maturation into the fully glycosylated receptor, whereas DOR maturation was significantly reduced in CNIH1-deficient cells at 6 h and 24 h (**Fig. 3f**). The findings support a role for CNIH1 in promoting DOR delivery to the Golgi apparatus, where terminal glycan processing occurs.

Given the impact on DOR surface levels, we next asked whether CNIH1 depletion affects DOR signaling. To assess G protein coupling upon receptor stimulation, we monitored miniGi recruitment to agonist-activated DOR using a split NanoLuc complementation assay^41^ in DOR-RUSH cells. After 1.5 h or 24 h of biotin treatment, DOR activation by the peptide agonist DADLE drove robust miniGi coupling in control cells, whereas CNIH1-deficient cells showed significantly reduced recruitment (**Fig. 3 g,h)**. As a second readout of signaling, we examined ERK1/2 phosphorylation downstream of DOR activation. After 24 h biotin treatment to drive receptor surface localization, stimulation of DOR-RUSH cells with the peptide agonist DPDPE triggered strong ERK1/2 phosphorylation in control cells but a markedly reduced response in CNIH1 KO cells (**Fig. 3i**). The results demonstrate that the diminished DOR PM levels in the absence of CNIH1 result in reduced DOR signaling capacity.

Taken together, CNIH1 is required for efficient ER-to-Golgi transport of DOR, which gates the conversion into the fully mature DOR glycoprotein and signaling.

### CNIH1 localizes to ER exit sites and the Golgi compartment

Next, we determined the subcellular localization of CNIH1. In HeLa cells, staining with a CNIH1 antibody revealed a signal in the perinuclear region and in puncta distributed throughout the cytoplasm (**Fig. 4a**). The CNIH1 antibody was highly specific, as the signal dropped to background levels in CNIH1 KO cells (**Extended Data Fig. 5a,b**). Visualizing different cellular (sub)compartments by antibodies or fluorescent protein-tagged marker proteins showed that the perinuclear pool of CNIH1 strongly overlapped with the cis-medial Golgi protein Giantin (**Fig. 4a**). Many of the CNIH1 puncta co-localized with Sec23, a COPII coat protein enriched at ER-exit sites (ERES)^42^ (**Fig. 4b**). Quantification using Pearson’s correlation and Manders’ colocalization coefficients indicated a high degree of overlap between CNIH1 with the Golgi and ERES, but minimal colocalization with endosomes marked by EEA1 or LysoTracker-labeled lysosomes (**Fig. 4c, Extended Data Fig. 5c**). A similar distribution of endogenous CNIH1 was observed in two additional cell lines, HEK293 and SH-SY5Y neuroblastoma cells **(Extended Data Fig. 5d)**. The results showed that CNIH1 localizes in the early secretory pathway, in particular ERES and the cis-medial Golgi, which are sites of COPII vesicles that transport transmembrane cargoes including GPCRs^8^.

**Figure 4:**
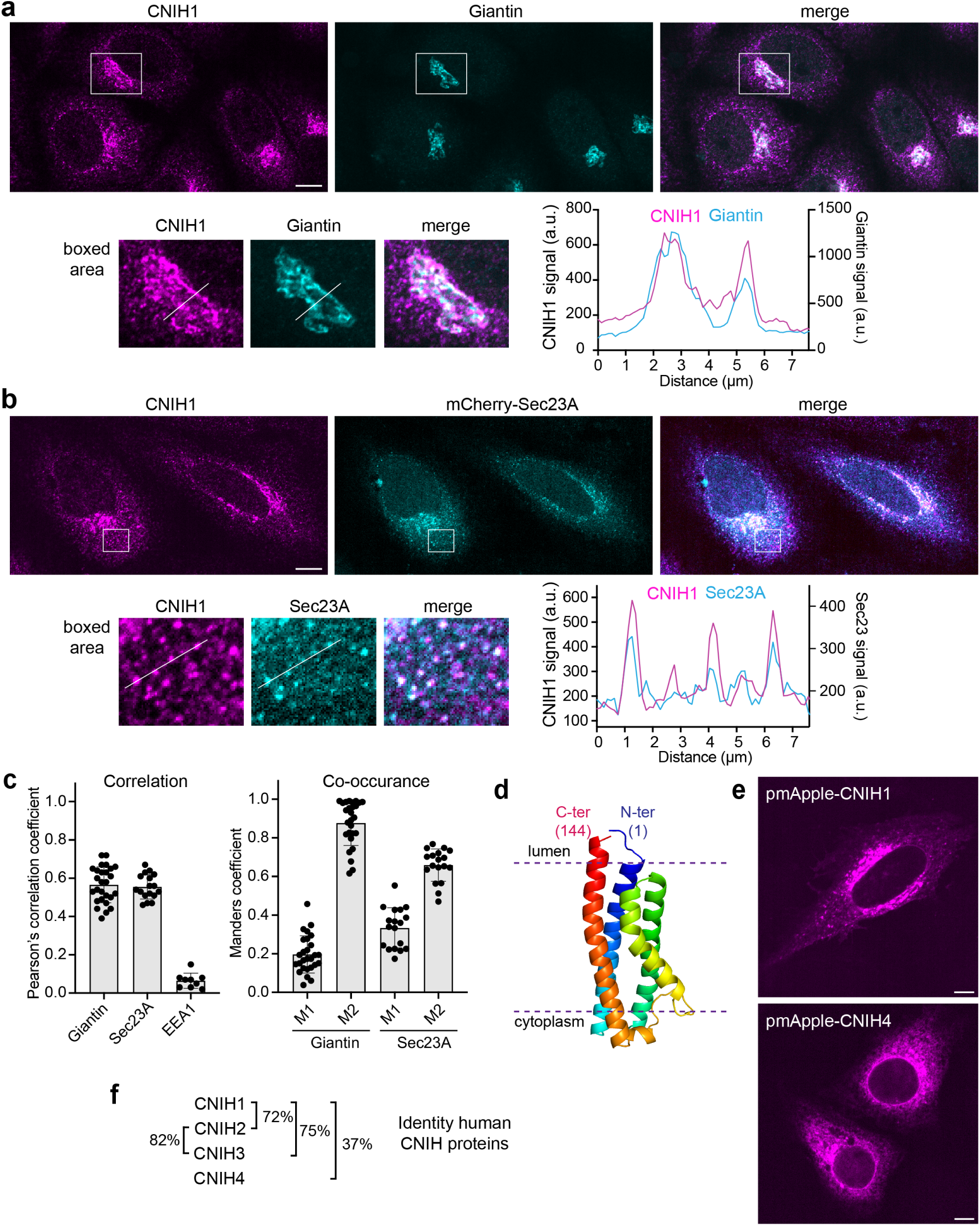
Subcellular localization of CNIH1 in ER exit sites and cis-medial Golgi. **(a)** Confocal microscopy image of HeLa cells immunostained for endogenous CNIH1 (magenta) and the cis-medial Golgi marker Giantin (cyan). Images are single confocal slices. The merged image shows strong co-localization in the Golgi region. The lower panel shows a magnified view of the boxed area. Line-scan analysis (right) displays the pixel intensity profiles of CNIH1 and Giantin along the indicated line. Representative image from n = 3 independent experiments. Scale bar, 10 μm. **(b)** Confocal microscopy image of HeLa cells immunostained for endogenous CNIH1 (magenta) and expressing the ERES marker mCherry-Sec23A (cyan). Images are single confocal slices. The merged image shows co-localization in the Golgi area and in puncta in the cytosol. The lower panel shows a magnified view of the boxed area, with a line-scan analysis (right) depicting pixel intensity profiles of CNIH1 and Sec23A across puncta at indicated line. Scale bar, 10 μm. **(c)** Left: Pearson’s correlation coefficient analysis to determine the degree of linear relationship between fluorescence intensities of CNIH1 and Giantin, Sec23A, or EEA1 in immunostained HeLa cells. Right: Manders’ colocalization coefficients M1 and M2 indicating overlap of CNIH1-positive pixels with Giantin or Sec23A (M1) or overlap of Giantin or Sec23A-positive pixels with CNIH1 (M2). The majority of Giantin- and Sec23A-positive pixels overlap with CNIH1. Cells analyzed: Giantin, n = 27; Sec23A, n = 17; EEA1, n = 9. **(d)** AlphaFold-predicted structural model of human CNIH1 (144 amino acids). The model has an average per-residue local distance difference test (pLDDT) confidence score of 84.3 and predicts a four-transmembrane-domain topology with the N-terminus (blue) and C-terminus (red) oriented toward the organelle lumen. The dotted line depicts the approximate membrane boundaries. **(e)** Confocal images of HeLa cells expressing CNIH1 (top) or CNIH4 (bottom), N-terminally fused to pmApple. Images are single confocal slices. Scale bar, 10 μm. **(f)** Percent identity between CNIH1 and other human CNIH paralogs. CNIH2 and CNIH3 show the highest identity.

CNIH1 is a multi-pass membrane protein with unknown structure. Yet, based on the high resolution structures of the highly homologous CNIH2 or CNIH3 proteins in complex with AMPA receptors^43,44^, and according to Alphafold prediction^45^, CNIH1 is expected to adopt a four-transmembrane domain topology with both N- and C-termini facing the organelle lumen, and the majority of the protein embedded in the membrane (**Fig. 4d**). When we expressed epitope- or fluorescent protein-tagged CNIH1 in HeLa cells, we detected a subcellular distribution that resembled endogenous CNIH1 (**Fig. 4e, Extended Data Fig. 6a**), with an ER and Golgi localization as described previously for transiently expressed CNIH1^46^.

The cornichon homolog protein CNIH4, previously identified to promote the surface expression of two distinct GPCRs^11^, exhibits 37% sequence identity with CNIH1 (**Fig. 4f**). CNIH4 did not score as a hit in our DOR-RUSH CRISPR screen; however, we extended our analyses to CNIH4 because of its homology to CNIH1 and its expression pattern, which parallels CNIH1 albeit at lower levels in most human tissues as well as in nociceptor sensory neurons that endogenously express DOR **(Extended Data Fig. 6b,c**)^47,48^. Epitope- or fluorescent protein-tagged CNIH4 localized more prominently to the ER when compared with CNIH1 (**Fig. 4e, Extended Data Fig. 6a**). Because both proteins are present in the early secretory pathway, we next tested the individual impact of CNIH1 and CNIH4 on surface targeting of diverse GPCRs as well as a non-GPCR cargo.

### CNIH1 selectively promotes the anterograde transport of a subset of GPCRs

We next tested whether CNIH1, identified here as a DOR export factor, also regulates the surface delivery of other secretory cargoes. To test this, we analyzed six class A GPCRs with distinct features: i) one highly homologous to DOR (MOR), ii) members of divergent subfamilies with different ligand types - muscarinic acetylcholine receptor M5 (M5R), histamine H1 receptor (H1R), purinergic P2Y1 receptor (P2YR), C-C chemokine receptor type 3 (CCR3) and orphan G-protein coupled receptor 35 (GPR35) - and iii) receptors with large extracellular (CCR3) or intracellular (M5R, H1R) domains relative to DOR. As a non-GPCR control, we included a glycosylphosphatidylinositol (GPI)-anchored GFP (GFP-GPI), a well-characterized model cargo that requires p24 cargo receptors for efficient export^49^. To examine biosynthetic transport, we generated stable RUSH HeLa cells for each protein and sorted cells for expression levels similar to the DOR-RUSH cells **(Extended Data Fig. 7a)**.

We then determined export kinetics of the GPCRs and GFP-GPI following biotin addition using flow cytometry to quantify surface levels. CCR3, M5R, H1R, and the GFP-GPI exhibited rapid ER-to-PM export, reaching maximal cell surface levels within 3 h of biotin-induced release (**Fig. 5a**), similar to MOR (**Fig. 1d**). In contrast, P2Y1 and GPR35 displayed markedly slower export kinetics (**Fig. 5a**), resembling DOR (**Fig. 1d**). The results showed that class A GPCRs differ in their ER-to-PM trafficking kinetics, suggesting cargo-specific regulation within the biosynthetic pathway. Next, we asked whether CNIH1 regulates the anterograde transport of the various cargoes. Comparing control and CNIH1-depleted cells, we found that CNIH1 knockdown significantly reduced surface targeting of MOR, CCR3, P2Y1, and GPR35, consistent with the effect on DOR (**Fig. 5b**). In contrast, PM transport of H1R, M5R, and GFP-GPI was unaffected. We then depleted CNIH4 by siRNA and performed the same assays. Unlike CNIH1 depletion, CNIH4 knockdown significantly reduced surface delivery of all tested GPCRs as well as GFP-GPI (**Fig. 5c**). The findings are consistent with a broad role of CNIH4 in secretory cargo transport, and a function for CNIH1 as a selective regulator for a subset of GPCRs. The results also imply that CNIH1 and CNIH4 have non-redundant cellular roles.

**Figure 5.**
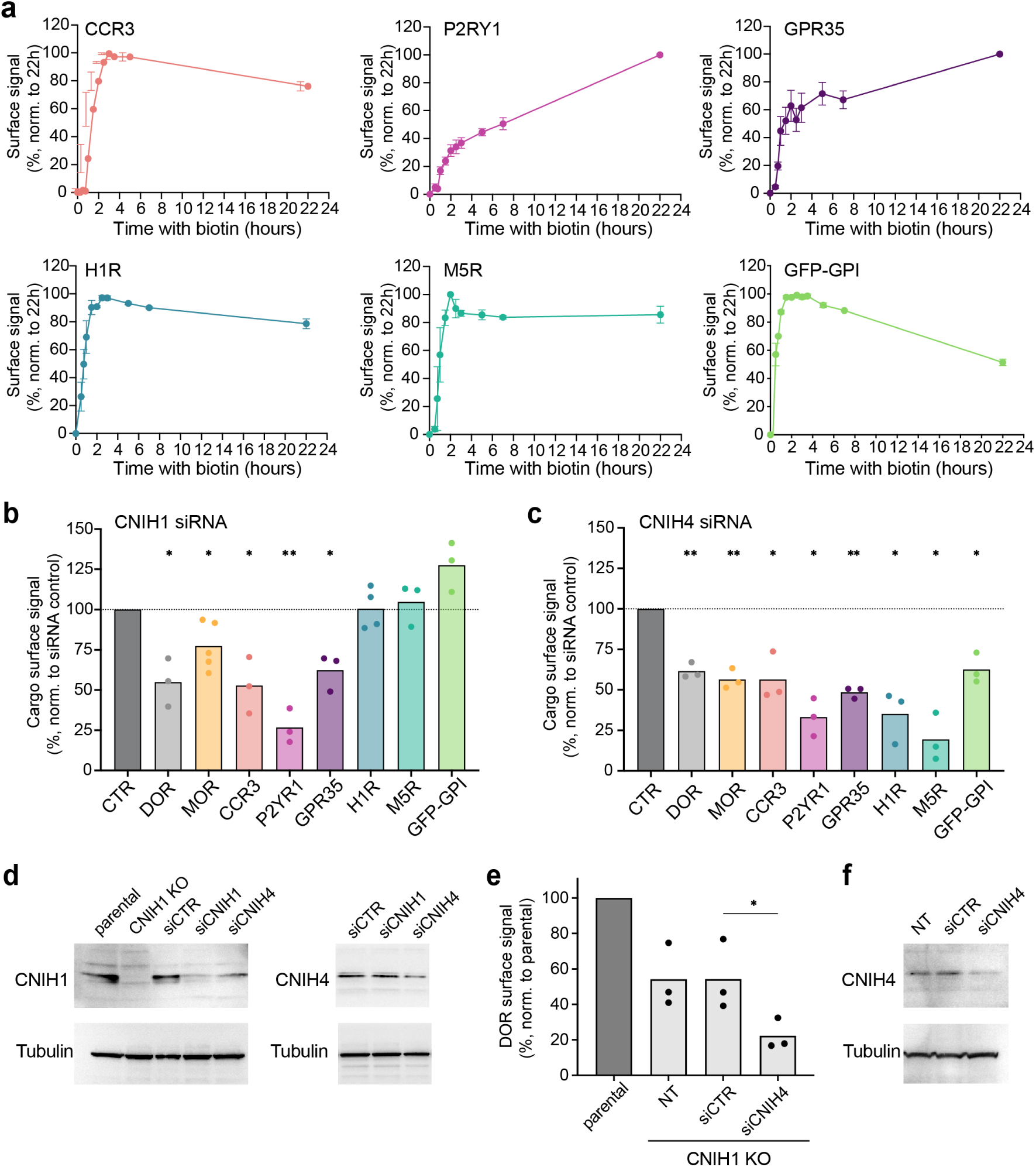
Selective regulation of GPCRs and non-GPCR cargoes by CNIH1 and CNIH4. **(a)** Kinetics of PM arrival of CCR3, P2YR1, GPR35, H1R, M5R and GFP-GPI in transgenic cargo-RUSH HeLa cells, measured by flow cytometry. The AF647 cargo surface signals at each time point were normalized to the condition with the highest surface levels (set to 100%) for each receptor. Data are mean ± SD from 3 independent experiments. **(b, c)** Effects of siRNA-mediated depletion of CNIH1 (b) and CNIH4 (c) on cell surface delivery of secretory cargos 1.5 hours after biotin addition. For each respective RUSH HeLa cell line, cells treated with non-targeting control siRNA (siCTR) served as a control (set to 100%). n ≥ 3 independent experiments. Significance was determined by one sample t-test. *, P < 0.05; **, P < 0.01. **(d)** Western blot analysis of CNIH1 (left) and CNIH4 (right) protein levels in HeLa cells treated with control siRNA (siCTR) or siRNAs targeting CNIH1 (siCNIH1) or CNIH4 (siCNIH4). Tubulin levels serve as a loading control. **(e)** DOR PM levels in DOR-RUSH CNIH1 KO cells further depleted of CNIH4 by siRNA after 1.5 h of biotin incubation. Signals were normalized to DOR-RUSH parental cells at 1.5 h biotin addition (set to 100%). Significance was determined by paired t-test. *, P < 0.05. **(f)** Western blot analysis of CNIH4 levels in DOR-RUSH CNIH1 KO treated with control siRNA (siCTR) or siRNA targeting CNIH4 (siCNIH4). Tubulin levels serve as a loading control.

Western blot analyses of CNIH1 and CNIH4 protein levels in siRNA-treated cells revealed that CNIH4 depletion led to the concomitant reduction in CNIH1 levels (**Fig. 5d, Extended Data Fig. 7b**). This suggested that CNIH4 stabilizes CNIH1 and showed that the inhibition of cargo export detected upon CNIH4 depletion was in fact the result of the combined reduction of CNIH4 and CNIH1 levels. To decipher CNIH4’s specific contribution to GPCR export, we treated DOR-RUSH CNIH1 KO cells with CNIH4 siRNA before quantifying DOR’s surface transport. Indeed, CNIH4 depletion in these cells further decreased DOR export, leading to a pronounced 80% inhibition of DOR surface targeting (**Fig. 5e,f**).

Together, these results demonstrate that both CNIH1 and CNIH4 contribute to GPCR export, but CNIH4 supports a broader range of secretory cargoes, whereas CNIH1 acts as a selective regulator for a specific subset of GPCRs.

### CNIH1 interacts with DOR and MOR, and receptor export depends on distinct CNIH1 domains

We next tested whether CNIH1 regulates opioid receptors through a direct physical interaction. Because both CNIH1 and the receptor are predominantly membrane-embedded, any complex between them would likely be sensitive to detergent solubilization. To stabilize potential interactions, we performed DOR immunoprecipitation following incubation with the cell-permeable chemical crosslinker DSP. Using DOR-RUSH HeLa cells, we detected a clear CNIH1 band upon isolation of DOR with GFP-trap beads (**Fig. 6a**). Co-immunoprecipitation experiments performed at different time points after DOR release from the ER indicated a reduced interaction between DOR and CNIH1 at 24 hours (**Fig. 6a**) when DOR is mature and predominantly localized in the PM (**Fig. 1,3**), suggesting that the interaction occurs within biosynthetic organelles.

**Figure 6:**
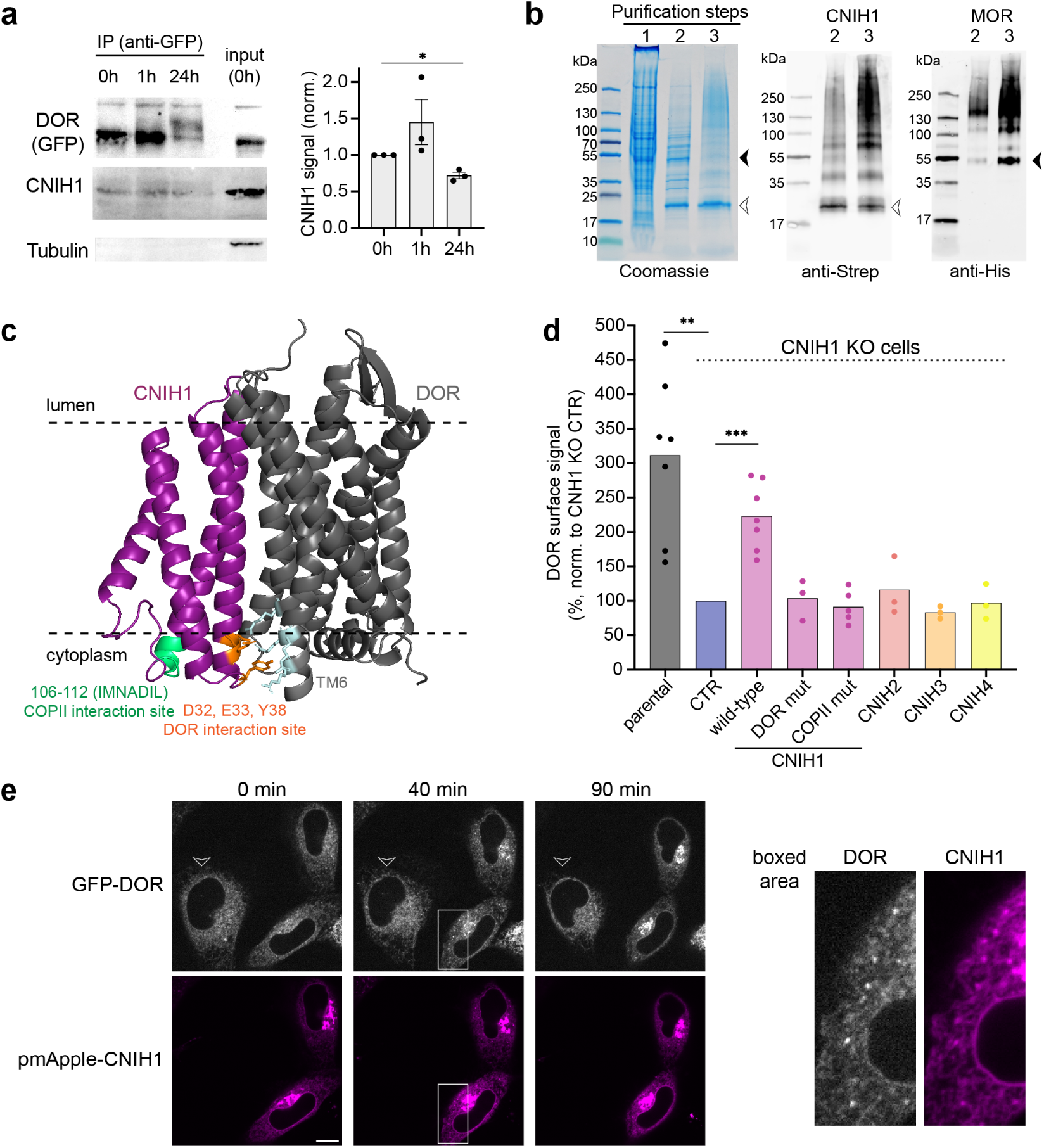
CNIH1 interaction with DOR and MOR, and identification of CNIH1 motifs required for DOR export. **(a)** Co-immunoprecipitation of DOR and CNIH1 in DOR-RUSH HeLa cells at 0, 1, and 24 h after biotin addition using GFP-Trap beads. Interaction was detected in the presence of the DSP crosslinker, tubulin serves as a negative control. Quantification of CNIH1 co-precipitation signals at different time points is shown normalized to the 0 h time point. Significance was determined by one sample t-test. *, P < 0.05. **(b)** Co-isolation of MOR and CNIH1 upon co-expression of His-tagged MOR and Strep-tagged CNIH1 in Sf9 insect cells. Sequential purification steps included: 1) Ni-NTA affinity purification of His-MOR; 2) StrepTactin affinity purification of Strep-CNIH1; and 3) size-exclusion chromatography. Elution fractions from each step were analyzed by SDS-PAGE and Coomassie stain or by western blot using anti-Strep and anti-His antibodies. Expected molecular weights of CNIH1 (open arrowhead) and MOR (filled arrowhead) are indicated. **(c)** Predicted structure of the CNIH1-DOR complex generated with Alphafold3 (interface predicted template modelling score, ipTM = 0.73). The interaction is predominantly predicted along DOR’s TMD6 and CNIH1’s TMD1, with a polar interface at the cytoplasmic side involving arginine residues in DOR (light blue) and aspartic acid (D32), glutamic acid (E33), and tyrosine (Y36) in CNIH1. The putative COPII binding site in CNIH1 (aa 106-112) is highlighted in green, with the sequence indicated. **(d)** Rescue of DOR export 1.5 h after biotin addition in DOR-RUSH CNIH1 KO cells by expressing CNIH1 constructs or human CNIH paralogs. CNIH1 KO cells were transfected with pm-Apple fused CNIH1 wild-type, CNIH1 DOR-interface mutant (D32A/ E33A/Y36A), CNIH1 COPII-binding mutant (aa 106-112 replaced by alanines), or CNIH2, CNIH3, CNIH4. Control (CTR) cells were transfected with pmApple. Parental DOR-RUSH cells transfected with pmApple serve as a control for DOR export at 1.5 h. n = 3-7 independent experiments. Significance was determined by one sample t-test. *, P < 0.05, **, P < 0.01. **(e)** Confocal time-lapse images of DOR-RUSH CNIH1 KO HeLa cells expressing or lacking (arrowhead) pmApple-CNIH1, acquired at the indicated time points after biotin addition (80 μM). Images are single confocal slices. The right panel presents an enlarged view of the boxed region, highlighting co-localized puncta. Scale bar, 10 μm.

As an orthogonal approach to probe the interaction of CNIH1 and opioid receptors, we co-expressed His-tagged MOR and Strep-tagged CNIH1 in baculovirus-infected Sf9 cells, an experimental system we have previously used to purify MOR in high yield and quality^50^. Following cell lysis and solubilization using mild detergents, we performed a three-step purification consisting of (i) Ni-NTA affinity capture of His-MOR, (ii) Strep-Tactin binding of Strep-CNIH1, and (iii) size-exclusion chromatography. Throughout the purification, protein complexity was monitored by SDS-PAGE and Coomassie staining, and the presence of MOR and CNIH1 examined by western blotting (**Fig. 6b**). Indeed, both CNIH1 and MOR were strongly retained in the final two purification elutions, appearing as clear bands at their expected molecular weights, along with higher molecular weight bands that point to the existence of higher order complexes. In sum, both purification strategies support the formation of a complex between CNIH1 and opioid receptors.

AlphaFold3 predictions suggested that the CNIH1 and DOR complex may form through polar interactions that involve Asp32, Glu33, and Tyr38 in the cytosolic end of CNIH1’s first transmembrane domain (TMD1) and arginines in DOR’s TMD6 (**Fig. 6c**). We also pinpointed a putative Sec 24/COPII interaction motif in CNIH1 at the end of the second cytoplasmic loop and TMD4 (106-IMNADIL-112) (**Fig. 6c),** based on insights into the function of the *S. cerevisiae* CNIH protein Erv14p^37^. We set out to test the functional role of CNIH1’s putative DOR and COPII interaction sites by performing rescue experiments in DOR-RUSH CNIH1 KO cells. First, we confirmed that reintroduction of wild-type CNIH1 by transient expression of pmApple-CNIH1 rescued efficient targeting of DOR to the cell surface at levels comparable with parental DOR-RUSH cells (**Fig. 6d, Extended Data Fig. 8a**). Live-cell imaging demonstrated enhanced Golgi arrival of DOR in cells expressing mApple-CNIH1, with colocalization in discrete cytosolic puncta (**Fig. 6e**), corroborating the role of CNIH1 in ER-to-Golgi export (**Fig. 3**). Next, we generated CNIH1 DOR and COPII interaction mutants, bearing alanine substitutions in the respective motifs. The mutant proteins were expressed at levels similar to wild-type CNIH1, showing that the mutations did not impact overall CNIH1 stability **(Extended Data Fig. 8b,c**). Yet neither the CNIH1 DOR mutant nor the COPII interface mutant rescued DOR surface transport (**Fig. 6d**). Microscopy analyses showed that both mutants were localized in intracellular organelles similar to wild-type CNIH1, although with slightly altered localization patterns **(Extended Data Fig. 8c)**: the CNIH1 COPII mutant accumulated in cytosolic puncta, while the DOR interface mutant was partially detected at the PM in addition to the Golgi, indicating that the domains contribute to the CNIH1 function and proper subcellular localization. Finally, we tested whether the other three human CNIH proteins can compensate for CNIH1 loss to promote DOR export. Expressing pmApple-tagged CNIH2, CNIH3, or CNIH4 in the DOR-RUSH CNIH1 KO cells, at levels comparable to CNIH1 **(Extended Data Fig. 8d)**, failed to rescue DOR surface targeting (**Fig. 6d**). This established that CNIH proteins act non-redundantly and that CNIH1 is a unique GPCR export factor.

## Discussion

The composition of the PM is dynamically shaped by the delivery of newly synthesized proteins from the secretory pathway, a process that also directly affects GPCR abundance and signaling capacity. Whether GPCR transport is selectively gated by dedicated regulatory factors, beyond the canonical secretory machinery, has remained an open question. To address this, we established a quantitative, synchronized RUSH-based assay that revealed GPCR-specific delivery kinetics and sensitively detected perturbations in ER-to-PM transport. Coupling this assay with a genome-wide CRISPR KO screen for DOR export and retention factors, we uncovered a broad network of both known and previously uncharacterized GPCR trafficking regulators. The recovery of canonical regulators, including COPII coat proteins and Rab GTPases, validated the screening approach, while the identification of new export factors, such as CNIH1, GOSR2, and Ric1, provides a new molecular framework for selective biosynthetic GPCR regulation. Intriguingly, several GPCR family members, such as GPR19 and GPR161, scored as DOR export factors, suggesting that GPCRs themselves may function as modulators within the secretory pathway, a concept that parallels findings for the Golgi-localized orphan receptor GPRC5A^51^. Among the retention factors we identified proteins with roles in protein folding and degradation, processes that likely contribute to GPCR protein homeostasis. Together, the screen uncovered a genetic map in human cells that governs GPCR transit through the secretory pathway.

In particular, our study establishes CNIH1 as a previously unrecognized export factor that directs the anterograde transport of DOR and a subset of diverse class A GPCRs. CNIH1 functions at the early stages of the secretory pathway. Loss of CNIH1 delays DOR transit from the ER to the Golgi and prevents its maturation into the fully glycosylated form, leading to a sustained deficit in PM localization and signaling. CNIH1 localizes to ERES and the cis-medial Golgi and directly associates with opioid receptors, consistent with a cargo receptor role that bridges recognition of GPCR clients to the cytosolic COPII machinery. CNIH1 mutants lacking the putative receptor- and COPII-interacting motifs were unable to support DOR export, highlighting distinct structural interfaces that coordinate cargo engagement and vesicle incorporation.

The cornichon family has previously been implicated in the secretory transport of different transmembrane cargoes. Mammalian CNIH2 and CNIH3 act as AMPAR receptor auxiliary proteins, binding AMPA subunits in the secretory pathway and remaining associated at the synapse to modulate receptor gating^52,53^. In contrast, our findings reveal that CNIH1 acts as a classical cargo receptor, similar to yeast Erv14p and *Drosophila* Cornichon, which mediate ER-to-Golgi transport without accompanying their clients to the cell surface^34,35,54^. The profiling of GPCRs for CNIH1 dependency indicates client specificity: not all GPCRs require CNIH1 for export. Structure predictions suggest that CNIH1 recognizes a positively charged patch on the cytoplasmic side of DOR’s TMD6. Interestingly, high-resolution DOR structures have revealed that certain TMD6 residues contribute to a polar network that stabilizes the inactive receptor conformation^55,56^, implying that CNIH1 may engage folding intermediates of some GPCRs. Consistently, the two CNIH1-independent GPCRs H1R and M5R exhibit rapid ER-to-PM transport kinetics, indicative of efficient folding and maturation. Interestingly, both H1R and M5R also have particularly long intracellular loop 3 (ICL3) regions that harbor several di-acidic (E/DxD/E) motifs, which may directly recruit COPII components and bypass the need for a cargo receptor^57^.

CNIH1 is broadly expressed across human tissues and shares an overlapping distribution with its homolog CNIH4. Despite this similarity, CNIH1 and CNIH4 exhibit distinct client specificities. Both promote ER export of nascent GPCRs, yet only CNIH1 rescues DOR trafficking in cells devoid of CNIH1, underscoring non-redundant functions among cornichon paralogs. Notably, depletion of CNIH4 led to a concomitant loss of CNIH1 protein, suggesting co-regulation at the level of stability, akin to other cargo receptor families such as the p24 proteins^58,59^. We propose that mammalian CNIH proteins have evolved a molecular division of labor: The highly similar CNIH1-3 serve as specialized cargo receptors for distinct structural classes of membrane-spanning clients, while CNIH4 functions as a broader regulator of export homeostasis. This diversification may reflect an evolutionary adaptation to the increasing complexity of receptor signaling networks in metazoans^60^.

The requirement of CNIH1 for efficient receptor maturation and surface expression directly impacts DOR signaling capacity. The findings demonstrate that biosynthetic trafficking is a primary determinant of GPCR signaling, complementing the mechanisms of receptor internalization and recycling. By governing the GPCR abundance at functional locations, CNIH1 likely influences spatially-encoded signals in polarized and neuronal cells, where distinct receptor pools elicit divergent physiological outcomes^61,62^. The identification of CNIH1 therefore introduces a new level of molecular precision to the canonical ER export machinery and suggests that GPCR signaling fidelity is in part encoded by GPCR-specific export adaptors.

Looking ahead, several questions emerge. How does CNIH1 achieve selectivity toward specific GPCRs, and what structural determinants define its client spectrum? Does CNIH1 operate cooperatively with ER-resident chaperones or quality control factors to license receptor export? Finally, since several human diseases arise from GPCR misfolding or trafficking defects^63^, CNIH1 could represent a novel target for modulating receptor export.

In summary, this work identifies CNIH1 as a central organizer of GPCR anterograde trafficking, uncovering a selective export route that connects the secretory machinery to receptor signaling. This study establishes a conceptual framework for how cargo receptors diversify secretory traffic and, more broadly, how membrane trafficking selectivity sculpts the functional GPCR landscape at the cell surface.

## Acknowledgements

We thank Arthur Radoux-Mergault for valuable pilot experiments, Amit Kumar and Zoé Valbret for tool generation, Andreas Boland for experimental input and support for X.Z., and Damien Jullié for discussing initial ideas. We are grateful to the members of the Flow Cytometry and Bioimaging Core Facilities for their expert support, and to Hadrien Soldati and Hanna Schwaemmle for advice on the CRISPR screen workflow. We also thank all members of the Stoeber lab for constructive feedback and ideas. This work was supported by the Swiss National Science Foundation (PCEFP3_181282 and 320030-231368), and by the Novartis Foundation for Medical-Biological Research.

## Author contribution

K.A. generated constructs and cell lines and performed the CRISPR screen. K.A., P.D., and I.M.-A. carried out functional studies. X.Z. purified the MOR-CNIH1 complex. S.B. co-supervised the screen and analyzed the NGS data. K.A., P.D., and M.S. designed experiments. All authors contributed to data analysis. M.S. conceived and directed the project and wrote the manuscript with input from all authors.

## Materials and methods

### Cell culture and plasmids

HeLa cells (CRM-CCL-2, American Type Culture Collection (ATCC), female) and HEK293 cells (CRL-1573, ATCC, female) were cultured in Dulbecco’s modified Eagle’s medium (DMEM, Gibco) supplemented with 10% fetal bovine serum (FBS, Gibco). Undifferentiated SH-SY5Y cells were cultured in 50% MEM Eagle’s EBSS (Sigma-Aldrich, M4655) and 50% Ham’s F-12 nutrient mix (Thermo Fisher Scientific, 11765054), supplemented with 10% FBS and 1 mM L-glutamine (Life Technologies, 25030024) at 37 °C in 5% CO₂.

Transgenic HeLa cell lines stably co-expressing Str-KDEL and N-terminal IL-2 signal sequence (ss) SBP-EGFP-tagged GPCRs (human), including the MOR-DOR chimera (human), were generated by cloning a “chicken β-actin promoter, Str-KDEL_SBP-EGFP-GPCR, PGK promoter, WPRE, antibiotic resistance” cassette into a PiggyBac transposon plasmid (Addgene #84239) and cotransfecting 2.5 µg of the plasmid with 1 µg of PiggyBac transposase using 3 µL Lipofectamine 2000 in a 6-well plate. Transfected cells were selected with antibiotics (2 µg/mL puromycin or 4 µg/mL blasticidin) and GFP-positive cells were isolated by FACS to achieve expression levels comparable between individual cell lines.

The MOR-DOR chimera was generated by cloning MOR lacking its C-terminal tail directly upstream of the DOR C-terminal sequence. Str-KDEL_SBP-EGFP-GPI was generated by fusing EGFP with a C-terminal CD59 GPI anchor downstream of Str-KDEL. All cloning into the PiggyBac transposon plasmid was performed using In-Fusion cloning (Takara). N-terminally tagged CNIH1 constructs, including CNIH1 wild-type, the DOR-interface mutant (D32A/E33A/Y36A), and the COPII-binding mutant (amino acids 106–112 replaced by alanine), were generated by cloning synthesized CNIH1 DNA fragments (Invitrogen GeneArt Synthesis) separated by a 10 amino-acid (2xGGGGS) flexible linker, with an N-terminal signal peptide and pmApple- or HA-tag, into pcDNA3.1. The ss-pmApple-tagged CNIH2, CNIH3, and CNIH4 constructs were generated by replacing CNIH1 in the ss-pmApple-CNIH1 pcDNA3.1 vector with synthesized CNIH2, CNIH3, or CNIH4 DNA fragments using restriction enzyme cloning. Lyn11-HA-SmBit was generated by cloning a synthesized Lyn11-HA DNA fragment (Invitrogen GeneArt Synthesis) into a SmBiT-containing pcDNA3.1 vector via restriction enzyme cloning. All plasmid sequences will be accessible in the Yareta data repository. Published plasmids used in this study include LgBiT-mGi1^17^, mCherry-Sec23A (Addgene #166894), mEmerald-Sec23A (Addgene #166893), Venus-EEA1, and pmApple-N1 (Addgene #54567).

### Ligands

DPDPE [D-Pen2,5]-enkephalin hydrate (#E3888), DADLE [D-Ala2, N-Me-Phe4, Gly5-ol]-enkephalin acetate salt (#E7131), and naloxone (#N7758) were purchased from Sigma-Aldrich. Ketanserin tartrate (#0908) was purchased from Tocris.

### Live cell imaging-based RUSH assay

GPCR-RUSH cells were seeded on poly-L-lysine-coated 35 mm glass-bottom dishes (MatTek, P35G-1.5-14-C). 16 to 24 h after seeding, live cells were imaged in HBS imaging solution (Hepes-buffered saline containing 135 mM NaCl, 5 mM KCl, 0.4 mM MgCl₂, 1.8 mM CaCl₂, 20 mM Hepes, and 5 mM D-glucose, pH adjusted to 7.4). Export was monitored using a spinning disk confocal microscope (Nipkow, Zeiss) with a Plan Apo 63×/1.4 oil DICIII objective in a temperature- and CO₂-controlled environment (37 °C, 5% CO₂). Biotin (80 µM final concentration; Sigma-Aldrich, B4501) was added by bath application. Images of the same cells were acquired before and at 2.5 to 5 min intervals after biotin addition.

### Flow cytometry-based RUSH assay

GPCR-RUSH cells were seeded in 12-well plates. 4 to 6 h after seeding, cells were incubated with 80 µM biotin at 37 °C for the 22 h time point. On the following day, cells were incubated with biotin for the indicated time points. At the end of the biotin incubation, surface receptors were labeled for 15 min at 37 °C with anti-GFP nanobody conjugated to AF647 (1:4000, the anti-GFP nanobody^25^ was expressed, purified, and conjugated as previously described^64^. Cells were washed three times with PBS (without Ca²⁺ or Mg²⁺) and detached by incubation with PBS-EDTA at 37 °C. Cells were resuspended in FACS buffer (PBS-EDTA or HBS imaging solution) on ice. Between 5,000 and 10,000 cells were analyzed by flow cytometry using a Beckman Coulter CytoFLEX Flow Cytometer. Cells were gated for (i) singlets and (ii) live cells. Data were analyzed using FlowJo software (version 10.10) by quantifying mean AF647 or GFP intensities.

### Pooled genome-wide CRISPR knockout screen

Approximately 100 million DOR-RUSH HeLa cells were transduced with the human CRISPR knockout pooled lentivirus library Brunello (lentiCRISPR v2, Addgene #73179-LV) at a multiplicity of infection (MOI) of 0.3. The lentiCRISPR v2 library comprises SpCas9 and 76,441 sgRNAs targeting 19,114 human genes as well as control sequences. 48 h after transduction, cells were selected with 2 µg/mL puromycin and expanded over 5–7 days, maintaining a minimum of 30 million cells representing ∼500x coverage. Following selection, the flow cytometry-based RUSH assay was performed. Control non-transduced cells were incubated with 80 µM biotin for 1.5 h and analyzed using a Beckman Coulter MoFlo Astrios or BD FACSAria Fusion flow cytometer. Gates were set for the top 10% and bottom 10% AF647 signal for subsequent sorting of transduced cells. Approximately 30 million cells were recovered from each gated population of transduced cells. Additionally, 30 million transduced, unsorted cells were saved for downstream screen normalization. Sorted cells were frozen until processing. Genomic DNA was extracted from the two FACS-sorted populations and the unsorted population using the QIAamp DNA Mini Kit (Qiagen, #51304). For each sample, sgRNA sequences were amplified by PCR as previously described ^30^. Parallel PCRs were performed, each containing 10 μg genomic DNA, 10 μL 5 μM uniquely barcoded P7 primer (Integrated DNA Technologies), 0.5 μL 100 μM P5 stagger primer mix (Integrated DNA Technologies), 8 μL dNTPs (Clontech, #RR001A), 1.5 μL ExTaq polymerase (Clontech, #RR001A), and 1× ExTaq buffer (Clontech, # RR001A) in a total volume of 100 μL. P5 and P7 PCR primer sequences are provided in the Supplementary Information Table 1. PCR cycling conditions were: initial denaturation at 95 °C for 1 min; 24 cycles of 95 °C for 30 s (denaturation), 53 °C for 30 s (annealing), and 72 °C for 30 s (extension); followed by a final extension at 72 °C for 10 min. PCR products were verified on a 2% agarose gel and purified using the AMPure purification system (Beckman Coulter, 63880). The sgRNA libraries were analyzed by next-generation sequencing (NGS) on an Illumina NovaSeq 6000 platform (SE 100bp reads) with approximately 50 million reads per condition. Library quality was assessed using Qubit and TapeStation. Sequencing was performed by the iGE3 Genomics Platform of the University of Geneva. Raw sequencing reads were aligned to sgRNAs and target genes using the MAGeCK algorithm ^32^, and output data were visualized using ggplot2.

### siRNA knockdown

Cells were seeded in 12-well plates and, 24 h later, transfected with 20 nM of specific individual siRNAs (Thermo Fisher Scientific, Silencer Select) using 2.5 µL Lipofectamine RNAiMAX transfection reagent (Thermo Fisher Scientific, #13778075). All siRNA sequences are provided in the Supplementary Information Table 1. 72 h after transfection, cells were assayed for GPCR export or knockdown efficiency by measuring mRNA levels (RT-qPCR) or protein levels (western blot).

### Real-time quantitative PCR (RT-qPCR)

Total RNA was extracted, and genomic DNA removed using the RNeasy Plus Mini Kit (Qiagen, #74134). RNA concentrations were quantified using a NanoDrop spectrophotometer (Thermo Fisher Scientific). Reverse transcription was performed with the High-Capacity cDNA Reverse Transcription Kit (Applied Biosystems, #4368814).RT-qPCR was carried out using PowerUp SYBR Green reagent (Applied Biosystems, #A25741) on a QuantStudio 3 or QuantStudio 5 thermocycler (Applied Biosystems). Specificity of the amplified products was confirmed by melt curve analysis. Relative RNA expression levels were calculated using the ΔΔCt method. All qPCR primer pair sequences are provided in the Supplementary Information Table 1.

### Generation of CNIH1 KO cell line

DOR-RUSH cells were seeded in 6-well plates and, 24 h later, transfected with 0.8 µg each of three sgRNA/Cas9 plasmids (pSpCas9(BB)-2A-Puro (PX459), Addgene #62988) targeting CNIH1. sgRNA sequences and the oligonucleotide sequences cloned into the sgRNA scaffold are provided in the Supplementary Information Table 1. Transfection medium was replaced after 6 h. 24 h post-transfection, cells were selected with 2 µg/mL puromycin for 48 h. Depletion of CNIH1 was confirmed by western blot and immunostaining of fixed cells.

### Immunoprecipitation

HeLa DOR-RUSH or DOR-RUSH CNIH1 KO cells were plated and treated with 80 µM biotin for 0, 1, 2, 6, or 24 h. For immunoprecipitation (IP), approximately 1 to 1.5 million cells were lysed in RIPA buffer (1% Triton X-100, 0.5% sodium deoxycholate, 0.1% SDS, 150 mM NaCl, 50 mM Tris, pH 7.4, 2 mM EDTA) supplemented with protease and phosphatase inhibitors (Roche, #11836170001 and #4906845001). Lysates were cleared of nucleic acid with Benzonase (Millipore, #E1014) for 20 min on ice. Cleared lysates were diluted in 10 mM Tris-HCl, pH 7.5, 150 mM NaCl, 0.5 mM EDTA to a final volume of 500 µL, of which 50 µL was retained as the input fraction. For IPs, GFP-Trap agarose beads (ChromoTek, gtak20) were incubated with the diluted lysates for 1.5 h at 4 °C. Beads were sedimented at 2,500 × g for 5 min at 4 °C, washed three times with wash buffer (10 mM Tris-HCl, pH 7.5, 150 mM NaCl, 0.05% NP-40, 0.5 mM EDTA), and centrifuged at 2,500 × g for 5 min at 4 °C. Washed beads were resuspended in 4× NuPAGE LDS Sample Buffer (Life Technologies, NP0007) with dilution buffer and incubated at 95 °C for 5 min.Proteins were separated on Bolt Bis-Tris Plus Mini Protein Gels, 4–12% (Invitrogen, NW04122BOX) or Novex Tris-Glycine Mini Protein Gels, 16% (Invitrogen, XP00162BOX), and transferred to Immobilon®-P PVDF membranes (Millipore, IPVH85R). Membranes were blocked with 5% BSA in TBS containing 0.1% Tween-20 for 1 h at room temperature, then incubated overnight at 4 °C with primary antibodies: CNIH1 (Sigma-Aldrich, HPA002544, 1:150), GFP (ProteinTech, 50430-2-AP, 1:3000), and recombinant mouse IG2A anti-α-tubulin (Geneva Antibody Facility, ABCD_AA345, 1:1000). Membranes were washed three times for 10 min with TBS-T (1× TBS, 0.1% Tween-20) and incubated with HRP-conjugated secondary antibodies (Jackson ImmunoResearch, 705-035-003, 1:2500) for 1 h at room temperature. After three washes, protein bands were visualized using SuperSignal West Pico PLUS Chemiluminescent Substrate (Thermo Fisher Scientific, 34580) and imaged with the iBright CL1500 system (Invitrogen, A44114). Band intensities were quantified using Fiji (ImageJ). For IPs probing biosynthetic DOR maturation based on glycosylation, final IP samples were treated with Endoglycosidase H (MedChem Express, HY-E70131, 1:10) for 1.5 h at room temperature prior to western blot analysis. For co-IPs, prior to immunoprecipitation, cells were washed twice and incubated in 1× DPBS (Gibco, 14190094). Cells were then treated with 2 mM dithiobis(succinimidyl propionate) (DSP) crosslinker (MedChem Express, HY-118759) for 2 h on ice. The crosslinking reaction was quenched with 50 mM Tris, pH 7.4, after which cells were washed with ice-cold PBS and lysed as described above. Following the immunoprecipitation protocol, the final diluted and boiled samples were treated with 500 µM DL-dithiothreitol (Sigma-Aldrich, D5545) to reverse the DSP crosslinking.

### Split NanoLuc-based G protein recruitment assay

HeLa DOR-RUSH parental or CNIH1 KO cells were seeded in 6-well plates and transfected with 0.8 µg Lyn11-HA-SmBit and 0.3 µg LgBiT-mG1 using 3 µL Lipofectamine 2000 (Thermo Fisher Scientific). 16 to 24 h after transfection, cells were treated with 80 µM biotin or left untreated for 1.5 or 24 h. Cells were then plated into black, clear-bottom 384-well plates at 20,000 cells/well in FluoroBrite DMEM (Gibco, #A1896701) containing Nano-Glo substrate (Promega, N2012) and incubated for 45 min at 37 °C. Luminescence was recorded at 37 °C on an FDSS/µCELL kinetic plate imager (Hamamatsu) equipped with an integrated dispenser. Following a 3 min baseline acquisition, 10 µM DADLE was added to each well, and luminescence was recorded every 10 s for 7 min. Two technical replicates were measured per condition. Raw luminescence values were first normalized to the baseline signal of each well (average signal during the 3 min pre-agonist baseline). Baseline-normalized values were then normalized to the vehicle-treated control to account for plate- and treatment-specific effects. Area under the curve (AUC) was calculated from the normalized time course following agonist addition.

### ERK1/2 activation assay

For ERK analysis by western blot, DOR-RUSH parental and CNIH1 KO cells were treated with 80 µM biotin in serum-free medium for 24 h, followed by stimulation with 10 µM DPDPE for 5 min. Cells were lysed in RIPA buffer and cleared with Benzonase nuclease as described in the “Immunoprecipitation” section. Protein concentrations were determined using the Pierce BCA Protein Assay Kit (Thermo Fisher Scientific, #23225) and adjusted in 4× NuPAGE LDS Sample Buffer (Life Technologies, #NP0007) containing 100 mM dithiothreitol to achieve a final concentration of 0.7 mg/mL protein per sample, resulting in 10 µg of protein per well in 15 µL loading volume. Proteins were separated on Bolt Bis-Tris Plus Mini Protein Gels, 4-12% (Invitrogen, NW04125BOX), and transferred to Immobilon-P PVDF membranes (Millipore, IPVH85R). Membranes were blocked with 5% BSA in TBS with 0.1% Tween-20 for 1 h at room temperature, followed by overnight incubation at 4 °C with primary antibodies: Phospho-p44/42 MAPK (Erk1/2) (Thr202/Tyr204) (Cell Signaling Technology, #4370S) and total p44/42 MAPK (Erk1/2) (Cell Signaling Technology, #4696S), both at 1:2,500 dilution. Membranes were washed three times for 10 min with TBS-T and incubated for 1 h at room temperature with secondary antibodies: CF770 Goat Anti-Rabbit IgG (Biotium, #20078) and CF680 Goat Anti-Mouse IgG (Biotium, #20065). Membranes were imaged using an Odyssey M Imager (LI-COR) with acquisition in the 700 nm and 800 nm channels. Band intensities were quantified using Fiji (ImageJ). Phospho-p44/42 MAPK signals were normalized to total p44/42 MAPK, and fold change was calculated relative to the unstimulated condition. Spectra Multicolor Low Range Protein Ladder (Thermo Fisher Scientific, #26628) was used as a molecular weight marker.

### Western blot

For western blot analysis, parental or CNIH1 KO DOR-RUSH cells, or cells treated with siRNAs, were lysed in RIPA buffer and cleared with Benzonase nuclease (see “Immunoprecipitation”). Protein concentrations were determined using the Pierce™ BCA Protein Assay Kit and adjusted in 4× NuPAGE LDS Sample Buffer, with 20 µg of protein loaded per well. Proteins were separated on Novex Tris-Glycine Mini Protein Gels, 16% (Invitrogen, XP00162BOX), and transferred to Immobilon-P PVDF membranes (Millipore, IPVH85R). Membranes were blocked with 5% BSA in TBS containing 0.1% Tween-20 for 1 h at room temperature, followed by overnight incubation at 4 °C with primary antibodies: CNIH1 (Sigma-Aldrich, HPA002544, 1:150), CNIH4 (Thermo Fisher Scientific, #PA5-113469, 1:1000), or recombinant mouse IG2A anti-α-tubulin (Geneva Antibody Facility, ABCD_AA345, 1:1000). Following primary incubation, membranes were washed three times for 10 min with TBS-T and incubated for 1 h at room temperature with HRP-conjugated secondary antibodies (Jackson ImmunoResearch, 705-035-003, 1:2500). After three additional washes, protein bands were visualized using SuperSignal West Pico PLUS Chemiluminescent Substrate (Thermo Fisher Scientific, 34580) and imaged with the iBright CL1500 system (Invitrogen, A44114). Band intensities were quantified using Fiji (ImageJ), and CNIH1 signals were normalized to tubulin as a loading control.

### Immunofluorescence

HeLa wild-type, DOR-RUSH, DOR-RUSH CNIH1 KO, HEK293, or SH-SY5Y cells were seeded on 15 mm glass coverslips in 12-well plates. When indicated, cells were treated with 80 µM biotin for the specified duration. Cells were fixed with 4% formaldehyde (Sigma-Aldrich, #D2650) in PBS (Life Technologies, #14040091) and blocked/permeabilized with 2.5% BSA (Sigma-Aldrich, A7906) and 0.1% saponin (Millipore, 558255) in PBS for 1 h at room temperature. Primary antibody incubation was performed overnight at 4 °C in a humidified dark chamber with the following antibodies: CNIH1 (Sigma-Aldrich, HPA002544, 1:100), Giantin (Geneva Antibody Facility, ABCD_AA341, 1:100), and HA-Tag (6E2) (Cell Signaling Technology, 2367S, 1:200). Coverslips were washed 3 times for 5 min each with blocking solution, followed by incubation with secondary antibodies for 1 h at RT: Goat anti-Human IgG Alexa Fluor™ 647 (Thermo Fisher Scientific, A-21445, 1:1000), Goat anti-Mouse IgG Alexa Fluor™ 488 (Thermo Fisher Scientific, A-11001, 1:1000), and/or Goat anti-Rabbit IgG Alexa Fluor™ 568 (Thermo Fisher Scientific, A-11036, 1:1000). After secondary antibody incubation, nuclei were stained with DAPI (Sigma-Aldrich, D9542, 10 µg/mL) for 10 min. Cells were washed 3 additional times for 5 min each and mounted using ProLong Glass Antifade Mountant (Thermo Fisher Scientific, P36982). Imaging was performed on a spinning disk confocal microscope (Nipkow, Zeiss) using Alpha Plan Apochromat 40× and 100×/1.46 Oil DIC objectives.

### Immunofluorescence-based CNIH1 subcellular localization assay

WT HeLa cells or RUSH-DOR cells were seeded onto 15-mm glass coverslips in a 12-well plate and after 24 h transfected with 0.15 μg Venus-EEA1, 0.15 μg mCherry-Sec23A or 0.15 μg mEmerald-Sec23A using 1.5 μl of Lipofectamine 2000. 16 to 24 h after transfection, cells were fixed using 4% FA in PBS or stained with 75 nM LysoTracker (Red DND-99, Thermo Fisher Scientific, L7528) for 30 min at 37°C prior to fixation. Cells were then permeabilized and blocked with 0.1% Saponin and 1.5% BSA in PBS and incubated overnight at 4°C with primary antibodies: anti-CNIH1 antibody (1:100; Merck, HPA002544) and recombinant human anti-giantin (1:100; Geneva antibody facility, ABCD_AA341) in blocking solution. After 3 washes, cells were incubated with goat anti-rabbit IgG Alexa Fluor 647 (Thermo Fisher Scientific, A32733) and goat anti-human IgG Alexa Fluor 405 (1:1000 Thermo Fisher Scientific, #A48275) for 45 min at RT. Samples were mounted using ProLong Glass Antifade Mountant (Thermo Fisher Scientific, P36982) and imaged with a spinning disk confocal microscope (Nipkow, Zeiss) using an Alpha Plan Apochromat 100x / 1.46 Oil DIC objective. Stacks (0.063 μm / 1 pixel) with 13x 0.5 μm z-steps were acquired for all wavelengths. Images shown are a single confocal slice (6th or 7th stack) where CNIH1 structures are most prominent. Regions of interest (ROI) were drawn around the cell for the single confocal slice. Two analyses were performed to quantify colocalization. (i) Using the ImageJ Coloc 2 plugin, the CNIH1 channel and the subcellular marker channel were analyzed to determine the Pearson’s correlation coefficient in the ROI applied to all the color channels. The results have value between −1 and 1. (ii) The signal was cleared from the outside of the ROI for all channels. Using the ImageJ Just Another Colocalization Plugin (JACoP), the CNIH1 channel and the subcellular marker channel were analyzed to determine the Manders’ overlap coefficient (M1 and M2) on the cleared images, keeping thresholding consistent for each experiment. M1 is the proportion CNIH1 signal coincident with a given subcellular marker signal. M2 is defined conversely. The results have values between 0 and 1.

### DOR export rescue experiments

DOR-RUSH and DOR-RUSH CNIH1 KO cells were seeded in 12-well plates and, after 24 h, transfected with 0.45 μg pmApple-tagged CNIH1 wild-type, CNIH1 DOR-interface mutant (D32A/E33A/Y36A), CNIH1 COPII-binding mutant (amino acids 106 to 112 replaced by alanine), CNIH2, CNIH3, CNIH4, or 0.2 μg pmApple-N1 using 1.5 μL Lipofectamine 2000. The following day, cells were incubated with 80 µM biotin for 1.5 h. At the end of the biotin incubation, surface receptors were labeled for 15 min at 37 °C with anti-GFP nanobody conjugated to AF647 (1:4000). Cells were washed three times with PBS (without Ca²⁺ or Mg²⁺) and detached by incubation with PBS-EDTA (Versene, Gibco) at 37 °C. Cells were resuspended in FACS buffer (PBS-EDTA) on ice. Between 10,000 and 15,000 cells were analyzed by flow cytometry using a Beckman Coulter CytoFLEX Flow Cytometer. Cells were gated for (i) singlets, (ii) live cells, and (iii) pmApple expression. Data were analyzed with FlowJo software (version 10.10) by quantifying mean AF647 and GFP intensity of transfected cells. Conditions were normalized to control cells (DOR-RUSH CNIH1 KO cells transfected with pmApple only).

### Purification of MOR-CNIH1 complex

Protein purification was performed as described previously^50^. Briefly, the genes encoding MOR and CNIH1 were cloned into pF1 vectors and co-expressed in *Spodoptera frugiperda* (Sf9) cells. The final MOR construct contained an N-terminal signal sequence, a FLAG tag, and a C-terminal HRV-3C protease cleavage site followed by an 8×His tag. The CNIH1 construct contained an N-terminal signal sequence, a FLAG tag, and a TEV protease cleavage site followed by a Strep II tag. Expression of MOR and CNIH1 was achieved by infection of Sf9 cells at a density of 2 × 10⁶ cells/ml. Infected cells were cultured at 27 °C for 48 h and harvested by centrifugation. Cell pellets were lysed and homogenized in buffer containing 10 mM HEPES pH 7.5, 10 mM MgCl₂, 20 mM KCl, protease inhibitor cocktail, and a nonspecific nuclease (supernuclease), followed by centrifugation at 40,000 rpm for 1 h. The resulting pellet was resuspended and solubilized in buffer containing 25 mM HEPES pH 7.5, 100 mM NaCl, 1% LMNG, 0.1% CHS, 10% glycerol, protease inhibitor cocktail, and 2 mg/ml iodoacetamide, and incubated at 4 °C for 4 h to solubilize membrane proteins. Insoluble material was removed by ultracentrifugation at 40,000 rpm for 1 h. The supernatant, supplemented with 10 mM imidazole, was incubated with Ni-NTA resin (Thermo Fisher Scientific) for 2 h. The resin was washed with buffer A (25 mM HEPES pH 7.5, 100 mM NaCl, 0.01% LMNG, 0.001% CHS, and 10% glycerol) containing 30 mM imidazole, and bound proteins were eluted with buffer A containing 300 mM imidazole. The eluate was loaded onto a 5 ml Strep-Tactin Superflow cartridge (Qiagen), washed with buffer A, and eluted with buffer A supplemented with 2.5 mM desthiobiotin. The eluted protein was further purified by size-exclusion chromatography using a Superose 6 Increase 3.2/300 column (Cytiva) pre-equilibrated with 20 mM HEPES pH 7.5, 100 mM NaCl, 0.001% LMNG, and 0.0001% CHS. Fractions containing the MOR-CNIH1 complex were pooled, concentrated, flash-frozen, and stored at −80 °C. Purified proteins were separated by SDS-PAGE and transferred to a nitrocellulose membrane. Membranes were blocked in PBST containing 3% BSA at RT for 1 h, incubated overnight at 4 °C with primary antibodies (anti-His or anti-Strep; Sigma-Aldrich), washed 5 times with PBST, and incubated with secondary antibody (donkey anti-mouse IRDye 680RD; LI-COR) for 1 h at RT. After five additional PBST washes, membranes were imaged using an Odyssey system (LI-COR).

---

**Extended Data Figure 1.**
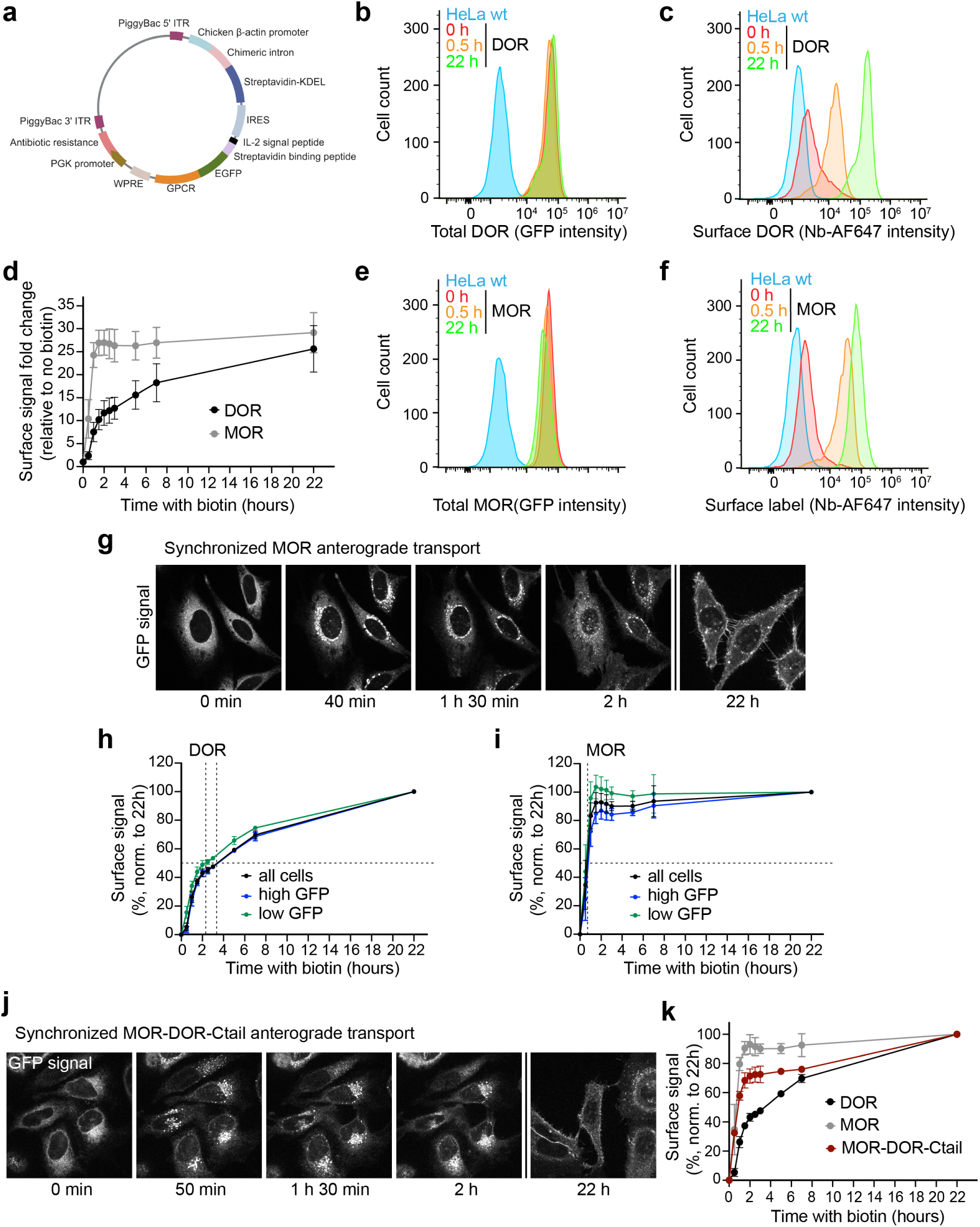
Characterization of DOR and MOR anterograde transport using the RUSH assay. **(a)** Schematic of the bicistronic GPCR-RUSH integrated via the PiggyBac transposon system. **(b, e)** Flow cytometry histograms showing total receptor expression based on GFP fluorescence in DOR-RUSH (b) or MOR-RUSH (e) cells, compared with HeLa wild-type (wt) cells. **(c, f)** Flow cytometry histograms of surface DOR (c) or MOR (f) levels at indicated time points after biotin addition. Surface receptors were stained with AF647-conjugated GFP-Nb. HeLa wt cells stained in parallel depicted as control. **(d)** Kinetics of DOR and MOR PM arrival, expressed as fold change in surface receptor levels relative to the signal at 0 h. Data are mean ± SD of 3 independent experiments. **(g)** Confocal time-series of MOR-RUSH HeLa cells imaged at the indicated time points after biotin addition (80 μM). At 22h, different cells are shown. Images are single confocal slices. Scale bar, 10 μm. **(h, i)** Kinetics of DOR (h) and MOR (i) PM arrival of cells with different receptor expression levels. The AF647 signal at each time point was normalized to the signal at 22 h (set to 100%). Data are related to Fig. 1d but include gating for the top 20% (high) or bottom 20% (low) expressing cells based on GFP. Half-times of export are indicated by dashed lines. Data are mean ± SD of 3 independent experiments. **(j)** Confocal time-series of MOR-DOR-RUSH HeLa cells at indicated time points after biotin addition (80 μM). At 22h, different cells are shown. Images are single confocal slices. Scale bar, 10 μm. **(k)** Kinetics of MOR-DOR PM arrival, compared to MOR and DOR transport. The AF647 signal at each time point was normalized to the signal at 22 h (set to 100%) for each receptor. Data are mean with ± SD of 3 independent experiments.

**Extended Data Figure 2.**
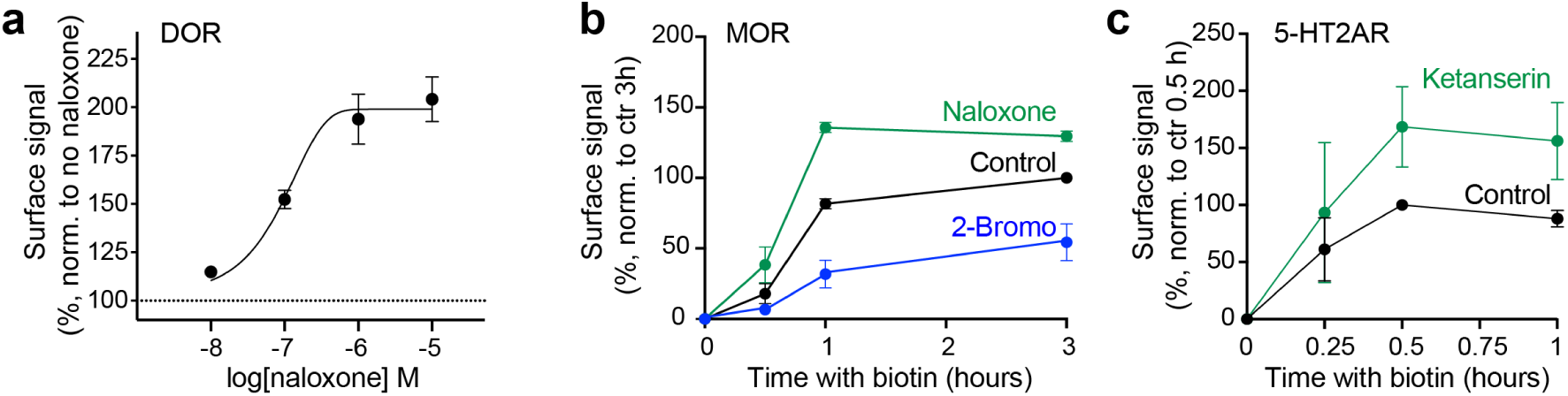
Characterization of regulated DOR, MOR, and 5-HT2AR anterograde transport. **(a)** Concentration-dependent effect of naloxone on DOR PM levels 1.5 h after biotin addition. AF647 signal at each naloxone concentration was normalized to the signal of the vehicle control (set to 100%). Data are mean ± SD of 3 independent experiments. **(b)** Kinetics of MOR PM arrival in cells pre-treated with naloxone (10 μM, 10 min pre-treatment) or 2-Br (100 μM, 16 h pre-treatment) prior to biotin addition. Signal at 3 h of untreated control cells was set to 100%. Data are mean +/- SD of 3 independent experiments. **(c)** Kinetics of 5-HT2AR PM arrival in 5-HT2AR-RUSH cells pretreated with 10 μM ketanserin, 10 min prior to biotin addition. Signal at 0.5 h of control cells was set to 100%. Data are mean +/- SD of 2 independent experiments.

**Extended Data Figure 3.**
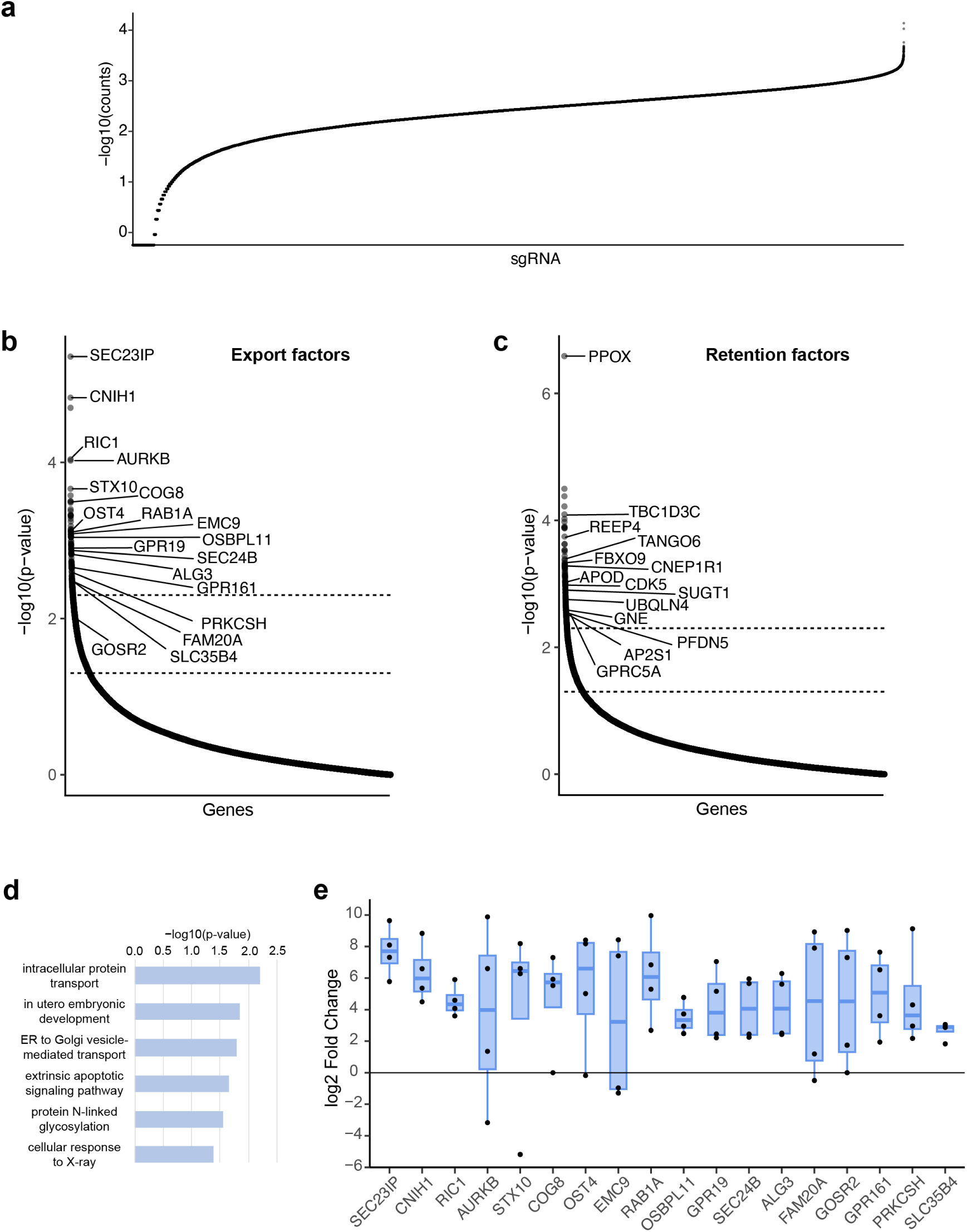
Genome-wide CRISPR/Cas9 screen identifies regulators of DOR anterograde transport. **(a)** Distribution of sgRNAs in unsorted control cells following transduction with the Brunello lentivirus library, demonstrating high diversity and coverage of sgRNAs. **(b, c)** Plot of export factors (b) or retention factors (c) from MAGeCK analysis, ranked by p-value. **(d)** Gene Ontology (GO) enrichment analysis of export factors using DAVID, showing functional categories significantly enriched among identified genes. Categories shown have p-value < 0.05. **(e)** Fold change in sgRNA counts between cell populations with high vs. low DOR surface levels. Highly ranked export factors are shown. Each data column indicates the fold change of the 4 individual sgRNAs targeting a single gene.

**Extended Data Figure 4.**
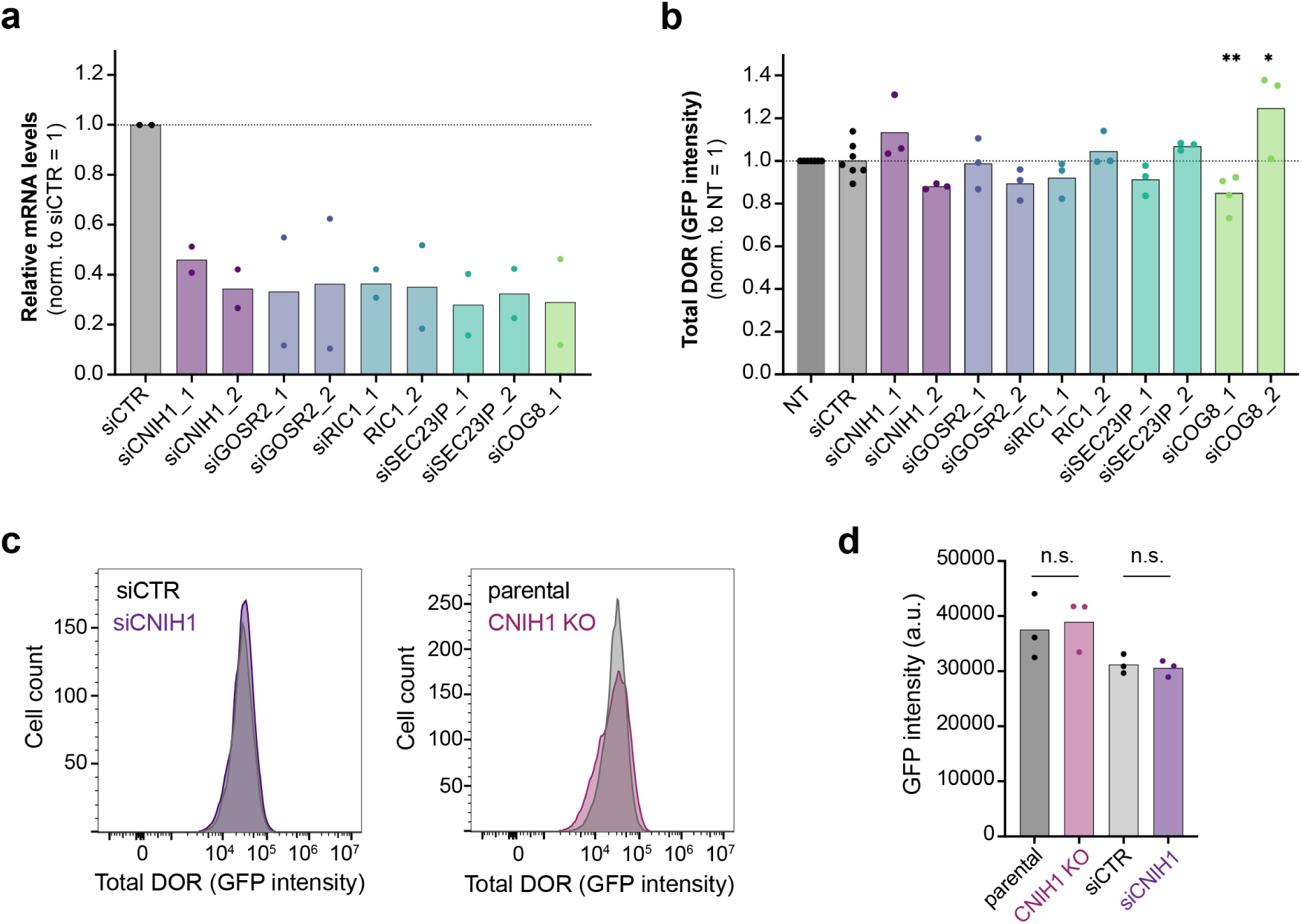
Validation of novel regulators of DOR anterograde transport. **(a)** Quantification of mRNA levels by RT-qPCR showing knockdown efficiency of target genes upon siRNA treatment. Expression was normalized to the non-targeting siRNA control (siCTR) set to 1 for each gene. n = 2 independent experiments. **(b)** Total receptor levels determined by GFP fluorescence intensity in DOR-RUSH HeLa cells treated with individual siRNAs as in (a). GFP intensity was measured by flow cytometry, and normalized to non-treated control cells (NT, set to 1). n = 3 independent experiments. Significance was determined by one-way Anova: *, P < 0.05; **, P < 0.01. **(c)** Flow cytometry histograms showing total receptor levels based on GFP fluorescence in DOR-RUSH cells lacking CNIH1. Left: Cells treated with control (siCTR) or CNIH1-targeting siRNA. Right: Parental and CNIH1 KO DOR-RUSH cells. **(d)** Quantification of total receptor levels based on mean GFP intensity in cells depleted of CNIH1 using siRNA or CRISPR/Cas9-mediated KO measured by flow cytometry. n = 3 independent experiments.

**Extended Data Figure 5.**
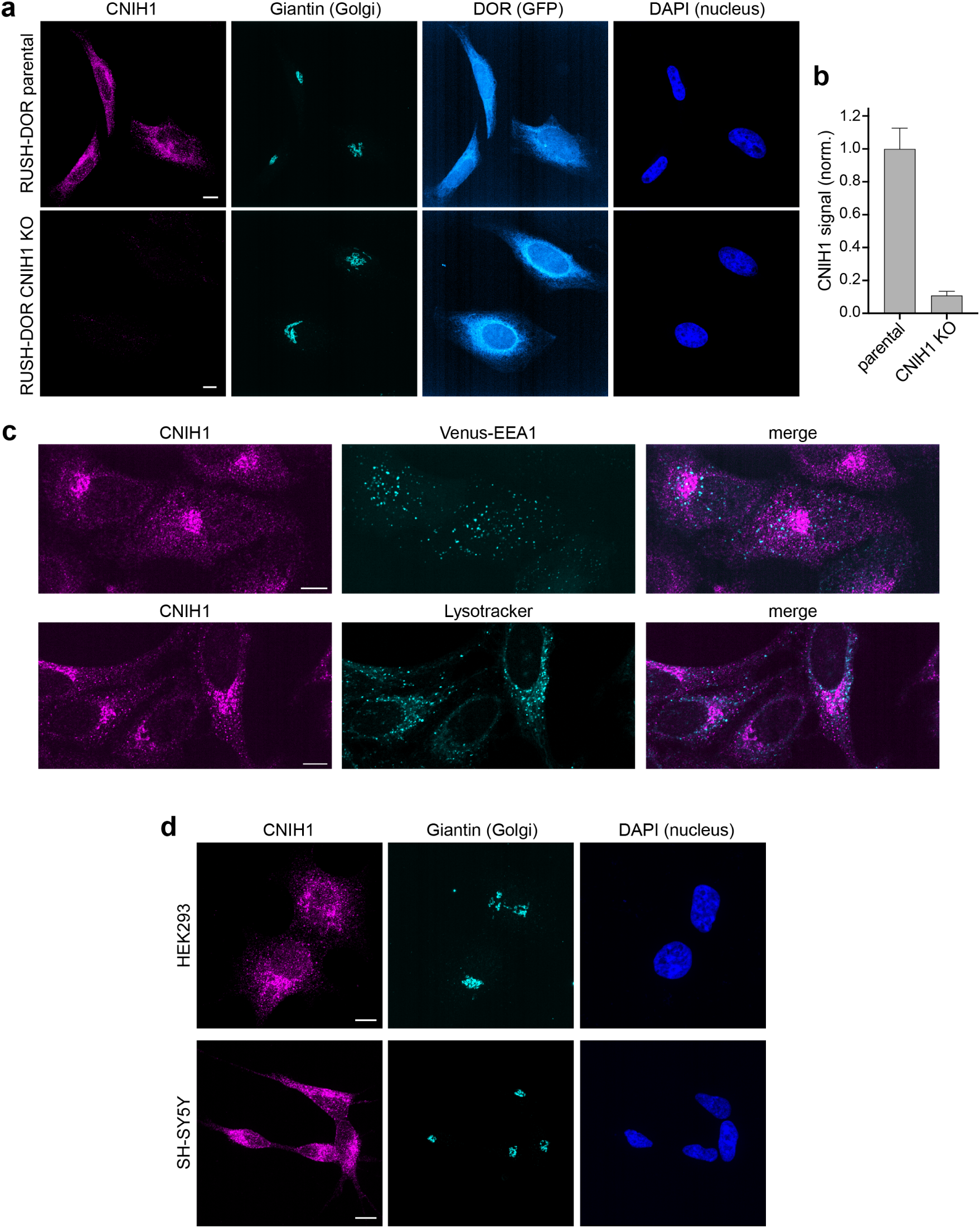
Subcellular localization of CNIH1. **(a)** Confocal microscopy images of parental (top) or CNIH1 KO (bottom) DOR-RUSH HeLa cells (no biotin added) immunostained for endogenous CNIH1 (magenta) and the cis-medial Golgi marker Giantin (cyan). DOR is shown by GFP fluorescence, nuclei were stained with DAPI. Images are maximum intensity Z-projections of confocal stacks. Scale bar, 10 μm. **(b)** Quantification of CNIH1 antibody staining intensity in the Golgi area in parental and CNIH1 KO DOR-RUSH HeLa cells. Data are mean ± SD, n = 4 independent experiments, each comprising 7-9 fields of view. **(c)** Confocal microscopy images of HeLa cells immunostained for endogenous CNIH1 (magenta) and either expressing the early endosomal marker Venus-EEA1 (cyan, top) or labeled with LysoTracker prior to fixation to visualize lysosomes (bottom). Merged images reveal no detectable co-localization between CNIH1 and endosomal or lysosomal compartments. Images are single confocal slices. Scale bar, 10 μm. **(d)** Confocal microscopy images of HEK293 (top) and SH-SY5Y (bottom) cells immunostained for endogenous CNIH1 (magenta) and the cis-medial Golgi marker Giantin (cyan). Nuclei were stained with DAPI. Images are maximum-intensity Z-projections of confocal stacks. Scale bar, 10 μm.

**Extended Data Figure 6.**
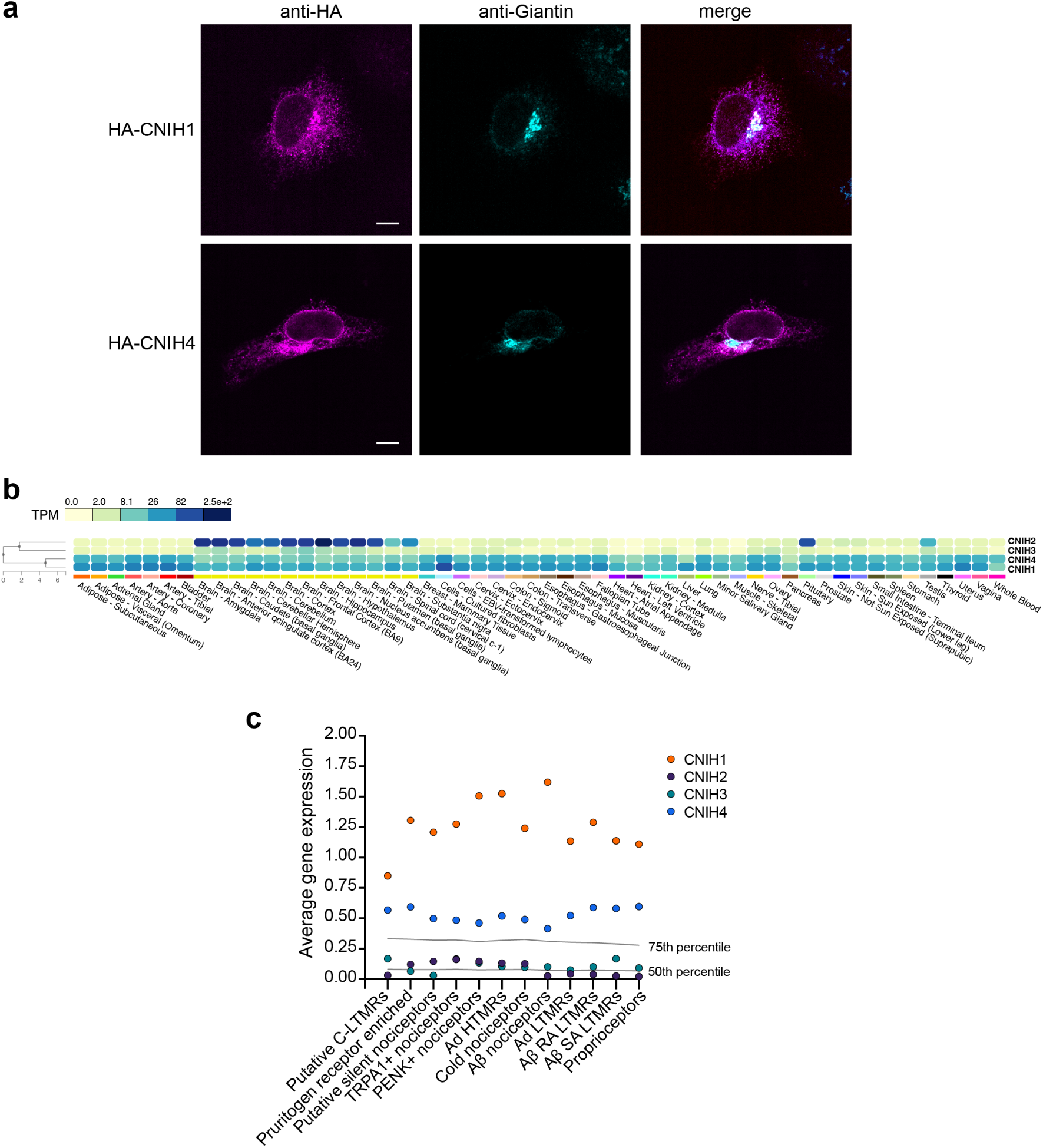
CNIH1 and CNIH4 subcellular localization and tissue or cellular expression patterns. **(a)** Confocal microscopy images of HeLa cells expressing CNIH1 (top) or CNIH4 (bottom) fused N-terminally to an HA tag. Cells were fixed and immunostained with anti-HA (magenta) and anti-Giantin (cyan) antibodies. Merged images show localization of CNIH1 in the Golgi area, which is less pronounced for CNIH4. Images are confocal slices. Scale bar 10 μm. **(b)** Heatmap of CNIH1-4 expression across diverse human tissues based on bulk RNA-sequencing data from the GTex portal (https://www.gtexportal.org/home/), using the Multi-Gene Query tool. Expression values are shown as Transcripts per Million (TPM). **(c)** Relative RNA expression levels of CNIH1-4 in a published dataset of 12 human nociceptor cell types (https://sensoryomics.shinyapps.io/RNA-Data/), known for endogenous DOR expression. Expression values of CNIH1-4 shown in colors indicated in the figure, with 75th and 25th percentile of overall RNA expression levels within each cell type shown as lines across the graph.

**Extended Data Figure 7.**
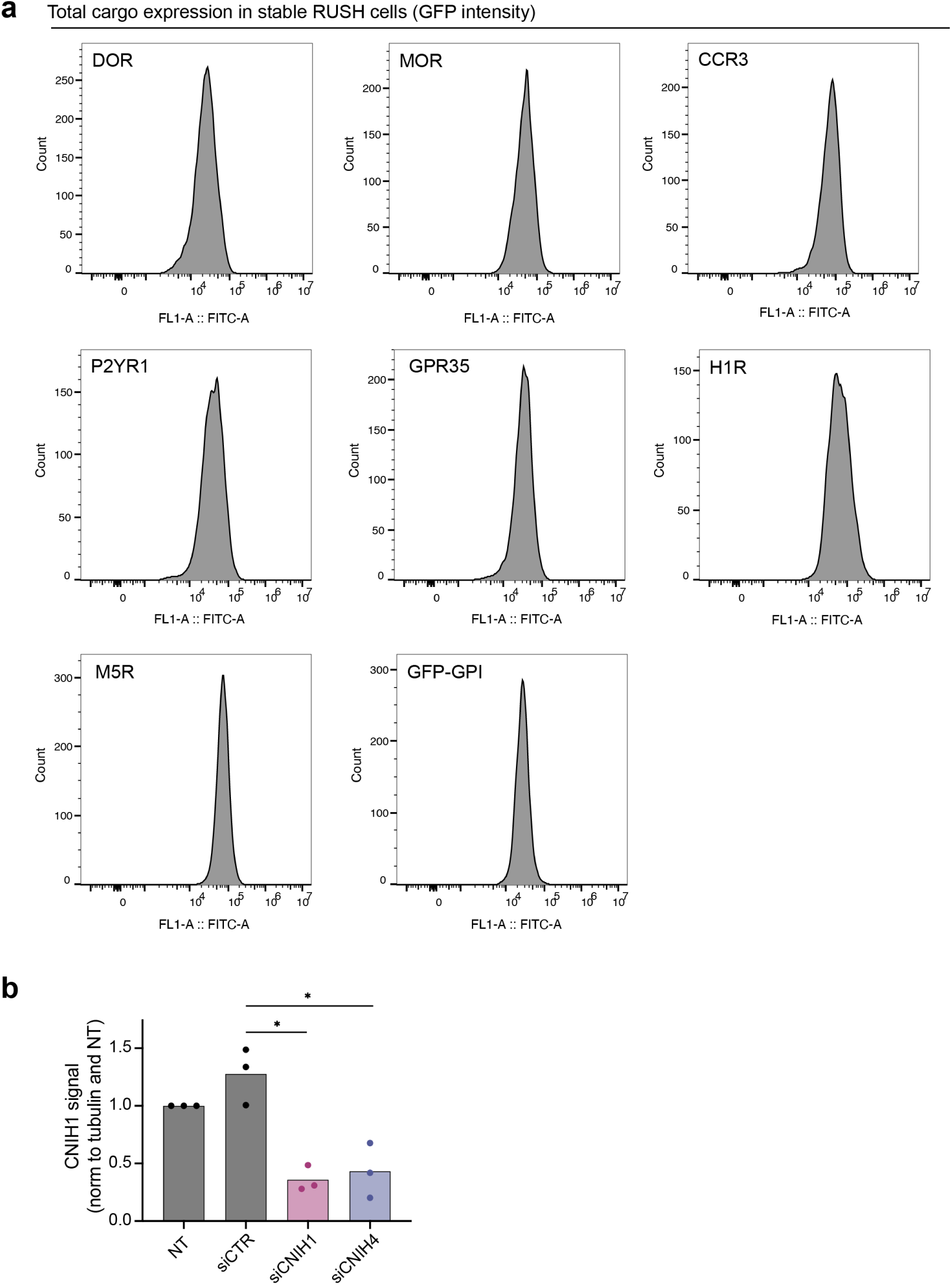
Characterization of anterograde transport using the RUSH assay after depletion of CNIH1 and CNIH4. **(a)** Flow cytometry histograms showing total cargo expression based on GFP fluorescence in various GPCR-RUSH and GFP-GPI-RUSH cell lines. Cargo expression levels are comparable to those observed for DOR-RUSH cells. **(b)** Quantification of CNIH1 protein levels in HeLa cells treated with control siRNA (siCTR) or siRNAs targeting CNIH1 (siCNIH1) or CNIH4 (siCNIH4), based on western blot analysis. CNIH1 signals were normalized to tubulin. n = 3 independent experiments. Statistical significance was determined by one-way ANOVA with Tukey’s multiple comparisons test. *, P < 0.05.

**Extended Data Figure 8.**
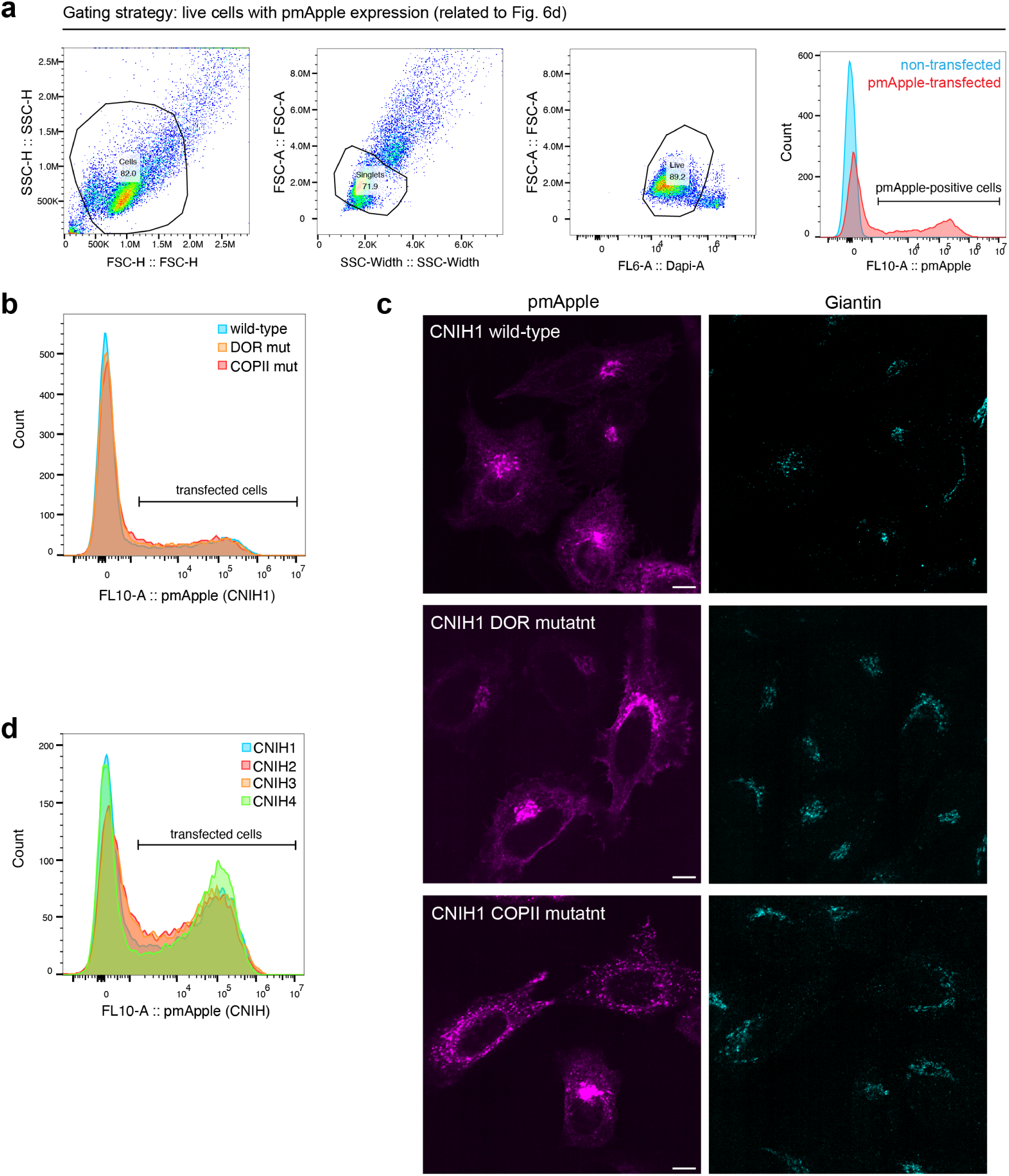
Flow cytometry and microscopy analyses of CNIH1 variants and CNIH paralogs. **(a)** Flow cytometry plots showing gating strategy for selecting pmApple-positive cells to assess effects on DOR export (related to Fig. 6d). Right: histograms of non-transfected and pmApple-N1-transfected cells used to define the pmApple-positive gate. **(b)** Representative flow cytometry histograms showing expression of pmApple-fused CNIH1 wild-type and DOR or COPII interface mutants in DOR-RUSH CNIH1 KO cells. Expression levels are comparable across constructs. **(c)** Confocal images of HeLa cells expressing CNIH1 wild-type or DOR and COPII interface mutants N-terminally fused to pmApple (magenta), fixed and immunostained for the cis-medial Golgi marker Giantin (cyan). Single confocal slices are shown. Scale bar, 10 µm. **(d)** Representative flow cytometry histograms showing expression levels of pmApple-fused CNIH1, CNIH2, CNIH3, or CNIH4 in DOR-RUSH CNIH1 KO cells. Expression levels are comparable across constructs.

